# Cohesin Acetylation and ATPase Activity Control Cohesion and Loop Architecture through Distinct Mechanisms

**DOI:** 10.1101/2025.10.30.685539

**Authors:** Lorenzo Costantino, Tiantian Ye, Kevin Boardman, Siheng Xiang, Jonathan Luo, Yudi Mu, Wenxiu Ma, Douglas Koshland

## Abstract

Cohesin is a conserved protein complex that mediates sister chromatid cohesion, chromosome condensation, gene regulation, and DNA repair. These processes rely on cohesin’s ability to tether sister chromatids and form chromatin loops, which depend on cohesin’s ATPase activity and Eco1-mediated acetylation of two lysine residues (K112 and K113 in budding yeast) in its Smc3 subunit. How cohesin’s ATPase activity and acetylation integrate to control cohesin functions remains poorly understood. Here, we analyzed chromatin architecture in yeast mutants with altered cohesin acetylation and/or ATPase activity. We find that acetylation of either K112 or K113 is sufficient to produce a wild-type chromosome structure with loops positioned at cohesin-associated regions (CARs), whereas loss of acetylation at both residues abolishes positioned loops, indicating that acetylation at either lysine alone can maintain wild-type chromatin architecture. We further show that a cohesin acetylation mutant, despite being defective in sister-chromatid tethering and thus failing to establish cohesion, still forms wild-type–like loops, while cohesion-competent mutants lack positioned loops. These results suggest that the activities required for cohesion and loop formation are mechanistically separable, arguing against passive loop capture. Moreover, a mutant with reduced ATPase activity showed a loop profile similar to wild type, indicating that cohesin with lower ATPase activity can still form wild-type chromatin architecture. By contrast, hyper-ATPase mutants accumulate positioned loops, suggesting that increasing ATPase activity can enhance loop processivity. Together, our findings support a multilayered regulatory model in which acetylation fine-tunes ATPase output and cohesin functions to shape genome architecture.

## Introduction

Cohesin is a conserved eukaryotic protein complex (Fig. 1A) that mediates sister chromatid cohesion, which is essential for chromosome segregation and cell viability. It also facilitates DNA repair, chromosome condensation, and regulates gene expression. Cohesin defects have been linked to birth defects and various diseases, including cancers (1). For many years, cohesin’s biological activities were believed to stem from its ability to tether two DNA regions within a chromosome or between sister chromatids (2, 3). However, recent in vitro studies have shown that cohesin and other members of the SMC (Structural Maintenance of Chromosomes) family of proteins can processively extrude DNA loops, a process named loop extrusion (4–6). This biochemical activity explains cohesin’s in vivo role in forming and maintaining dynamic, genome-wide intrachromosomal loops detected by chromosome conformation capture techniques (7– 11). Tethering and loop extrusion require cohesin’s ATPase activity and are regulated by Eco1 acetylation of cohesin’s Smc3 subunit (5, 6, 12–15). A deeper understanding of the roles and interplay of cohesin ATPase and acetylation in tethering and looping is crucial for comprehending cohesin’s capacity to promote diverse biological functions.

**Fig. 1.**
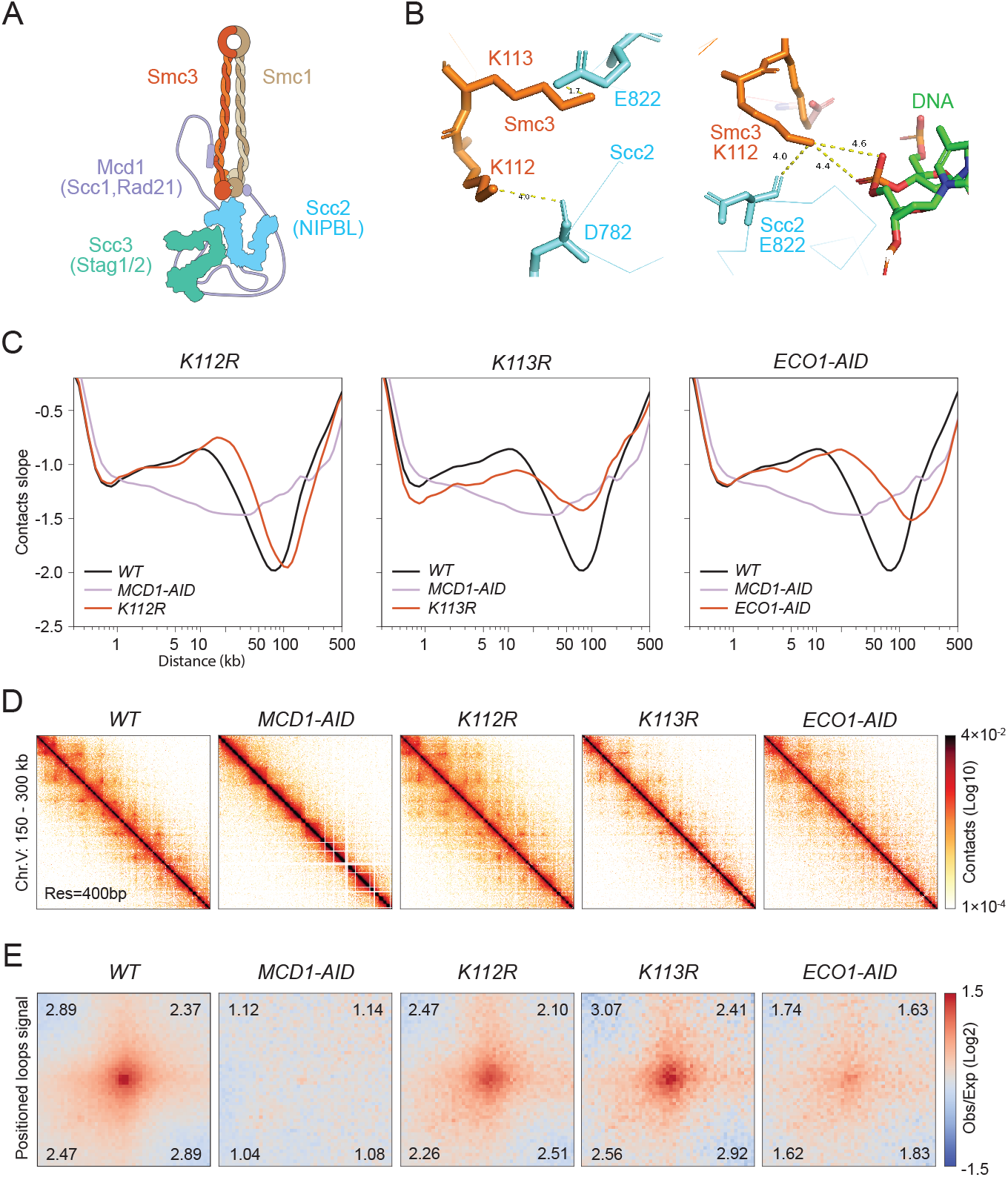
Chromosome structure in wild-type and cohesin acetylation mutants. **(A)** Cartoon depiction of the cohesin complex bound by the loader subunit Scc2p. Protein subunit names shown are from budding yeast, with alternative names for yeast (Scc1) and human orthologs (Rad21, NIPBL, Stag1/2) indicated in parentheses. **(B)** Cryo-EM structure of *S. cerevisiae* cohesin (PDB ID: 6ZZ6; Collier et al., 2020). The left panel highlights Smc3-K113 proximity to Scc2-E822; the right panel shows Smc3-K112 proximity to DNA and Scc2-D782. **(C)** Chromosome contacts in cohesin acetylation mutants. Micro-C XL analysis of chromosome interactions in mitotically arrested wild-type cells (*WT*), single acetylation-deficient mutants smc3-K112R (*K112R*) and smc3-K113R (*K113R*), and cells depleted of Eco1 acetyltransferase (*ECO1-AID*) or cohesin (*MCD1-AID*) (strains genotypes in Table 1). Derivative slopes of the contact frequency curves (shown in Fig. S1A) are plotted against genomic distance in black for *WT*, mauve for *MCD1-AID*, and red for acetylation-mutants. **(D)** Contact maps in cohesin acetylation mutants. Micro-C XL contact maps at 400bp resolution over an arm region at chromosome V 150-300kb for the indicated strains listed in C. Interaction intensity in contact maps will be represented throughout the paper using a standard log10 scale colormap, ranging from white (no detectable interactions) to black (strongest interactions). **(E)** Genome-wide signal for positioned loops at CARs in cohesin acetylation mutants. Piled-up heatmap of the ±5kb regions centered at CARs for the indicated strains listed in C. Numbers in the corners represent the fold-change of the signal enrichment of the center pixel over the indicated corner pixels. The pile-ups throughout the paper will utilize a diverging colormap for observed/expected signal in log2, with positive signal enrichment represented in red and negative signals in blue.

Cohesin ATPase and acetylation have been intensively studied, revealing a complex relationship. Cohesin forms two ABC-like ATPase active sites through dimerization of its Smc1 and Smc3 subunits (16). The ATP binding and ATPase activity of both active sites are required for cohesin binding to DNA, and hence for all cohesin’s functions, including cell viability (12, 13). Eco1 acetylation of cohesin is also essential for cell viability (14, 15, 17–19). It promotes stable DNA binding (20), the formation of cohesion by tethering the second sister chromatid (21), and the control of loop size (22, 23). Eco1p acetylates a conserved lysine in cohesin’s Smc3 subunit (K113 in budding yeast and K106 in mammals) (Fig. 1B) (14, 15, 19). Several studies have exploited K113 mutations that block (K113R) or mimic (K113Q) acetylation, suggesting that regulated K113 acetylation is essential for viability and cohesion (14, 15, 19). Follow-up studies indicated that K113 acetylation downregulates cohesin ATPase activity by perturbing its binding to, and therefore activation by, Scc2 (NIPBL in mammals) (Fig. 1B) (21, 24, 25). Scc2 is necessary for promoting both chromosome binding in vivo and loop extrusion in vitro (5, 6, 26).

Several important questions regarding cohesin’s ATPase and acetylation remain unanswered. For example, establishing the role of cohesin ATPase in DNA binding was critical. However, DNA binding is only the first step in generating cohesion or DNA loops. The contribution of cohesin ATP hydrolysis to subsequent steps, like capturing a second DNA strand or modulating loop size, is unknown. As a second example, Eco1p also acetylates a conserved lysine Smc3-K112 (K105 in mammals) that is adjacent to K113 (Fig. 1B) (14, 15, 19). The analysis of K112 acetyl-null (K112R) showed that K112 acetylation was not required for viability or cohesion. In the absence of a biological function, the roles of K112 acetylation in DNA binding, ATPase activity, or loop formation have not been investigated. The proximity of K112 to an aspartate on Scc2 and to a phosphate on the DNA backbone (Fig. 1B) suggests K112 might also promote Scc2 stimulation of cohesin ATPase in response to DNA binding. If so, does K112 acetylation impact loop size or position? To answer these and related questions, we studied the impact of a panel of hypomorphic mutations that alter cohesin ATPase activity or its acetylation on looping and tethering.

## Results

### K112 or K113 acetylation regulates chromatin loop size and positioning

Eco1p acetylation of cohesin is required to constrain loop size and form positioned loops, since depleting Eco1p or mutating both K112 and K113 to acetyl nulls results in loop expansion (22, 23). However, the individual contributions of K112 and K113 acetylation in looping or controlling ATPase levels have not been studied. To understand the role of K112 and K113 acetylation in chromosome structure, we utilized two acetyl-null mutants at these positions. These strains expressed only a Smc3 subunit that was either blocked for K112 acetylation (smc3-K112R) or K113 acetylation (smc3-K113R SMC3-AID in the presence of auxin). We compared the chromosome architecture in these strains by arresting them in mitosis and subsequently performing Micro-C XL (11, 27). As controls, we also used Micro-C XL to analyze chromosome architecture in Eco1-depleted (ECO1-AID in the presence of auxin) or cohesin-depleted (MCD1-AID in the presence of auxin) cells. For brevity and clarity, we referred to these strains in the remainder of the text and figure legends as K112R, K113R, ECO1-AID, and MCD1-AID, respectively. We also compared their structures to those found by Hi-C analysis of the smc3-K112R-K113R double mutant (K112R-K113R) and its corresponding wild type (22).

We began our Micro-C XL analysis by plotting contact decay curves, which show the average frequency of chromatin contacts as a function of genomic distance— that is, how often two genomic regions interact based on their separation (Fig. S1A). To more precisely compare strains, we also examined the derivative (slope) of these curves, which captures the rate of change in contact frequency across genomic distances (Fig. 1C). As previously shown (11), wild-type cells exhibited both elevated contact frequencies and more positive derivative slopes compared to cohesin-depleted cells, indicating that interactions around the 10 kb range are largely dependent on cohesin (WT vs. MCD1-AID in Fig. 1C and Fig. S1A). Interestingly, the K112R mutant closely resembled the wild-type positive slope with enriched contacts in the 20–50 kb range, suggesting good looping activity with slightly bigger loops being formed. The K113R mutant also displayed a clear gain in contact frequencies and positive slopes relative to cohesin-depleted cells, though somewhat attenuated compared to wild type. These profiles indicate that K113R retains substantial loop-forming capacity, suggesting that looping could occur independently of cohesin’s tethering activity, since the K113R mutant was defective in sister chromatid cohesion (14, 15).

In contrast, yeast cells expressing cohesin lacking acetylation on both K112 and K113 residues (ECO1-AID or K112R-K113R double mutant) exhibited a marked increase in both derivative slopes and contact frequencies in the 50–200 kb range compared to wild type, K112R, or K113R single mutant cells (Fig. 1C and Fig. S1A) (22, 23). This substantial gain in long-range interactions confirmed that Eco1-mediated acetylation limits the size of chromatin loops (22, 28). Notably, the wild-type loop profiles observed in the K112R or K113R single mutants imply that acetylation at either K113 or K112 is sufficient to limit the size of chromatin loops.

In wild-type cells, cohesin-dependent interactions detected by chromosome conformation capture techniques are thought to arise from the crosslinking of sequences, which can occur either within or between sister chromatids (29, 30). In contact maps, the diffuse off-diagonal “fuzz” reflects random looping events with variable anchor positions, whereas distinct off-diagonal “spots” represent positioned loops anchored at specific, recurrent genomic sites that correspond to cohesin-associated regions (CARs) in budding yeast (Fig. 1D and Fig. S1B). To quantify genome-wide the presence of positioned loops anchored at CARs, we performed a pile-up analysis of contact maps centered on CARs using 5 kb flanking regions, which revealed a genome-wide enrichment of signal above the background level of random distal interactions in wild type (Fig. 1E). Strikingly, these off-diagonal spots were also present in the contact maps and pile-ups of the K112R and K113R single mutants, with signal intensities comparable to those observed in wild-type cells (Fig. 1D and 1E; Fig. S1B), indicating that positioned loop formation is largely retained in these strains. In contrast, cells lacking both acetylations (ECO1-AID and K112R-K113R double mutant) showed a dramatic reduction in both the visibility of off-diagonal spots and the signal intensity in pile-ups (Fig. 1D and 1E; Fig. S1B) (22, 23). These results suggest that acetylation of either K112 or K113 is sufficient to support the formation of positioned loops, while simultaneous loss of both modifications severely impairs this process, resulting in loop expansion.

Positioned loop anchors in wild-type cells correlate with the crosslinking of two CARs, most prominently between adjacent ones (+1), followed by a gradual decrease with more distal CARs until the +5 CAR (Fig. S1C) (11). The K112R and K113R single mutants displayed a slightly increased intensity of distal interactions between CARs compared to wild type, with contacts extending to more distal CAR sites (Fig. S1C). In contrast, the distal interactions between CARs were significantly reduced in Eco1-depleted and K112R-K113R double mutant cells (Fig. S1C). Together, our results suggested that the accumulation of CAR-associated distal interactions required the acetylation of either K112 or K113.

We used chromatin immunoprecipitation (ChIP) to assess whether the differences in chromosome structure in the mutants correlated with cohesin binding to DNA. Aliquots from the same cultures were used for Micro-C XL and subjected to ChIP-seq to qualitatively evaluate the genome-wide pattern of cohesin binding, or ChIP-qPCR to quantitatively assess binding at representative CARs (Fig. S2A–C). ChIP-seq revealed a global pattern of cohesin binding at CARs that was indistinguishable between the mutants and the wild type (Fig. S2A). The qPCR analysis showed similar levels of cohesin binding at a representative CAR and at the centromere across all mutants (Fig. S2B). Thus, the level of cohesin binding to chromosomes could not explain the differences in chromosome structure in cells with cohesin mutant K112R (only K113 can be acetylated) and K113R (only K112 can be acetylated) compared to those with unacetylated K112 and K113 (ECO1-AID).

Recent studies suggest that Pds5 binding to chromatin-bound cohesin is responsible for limiting the length of distal interactions and the formation of positioned loops (22, 23). Furthermore, acetylated cohesin binds Pds5 better (22). Thus, we hypothesized that Pds5 binding to cohesin might be unaffected in the wild type or cells with singly acetylated cohesin (K112R or K113R), leading to similar distributions of distal interactions. In contrast, Pds5 binding might be compromised in cells with unacetylated cohesin (ECO1-AID), leading to longer distal interactions and the loss of positioned loop anchors.

To test this model, we used ChIP-qPCR to assess Pds5 binding at CARs on chromosome arms and centromeres. The Pds5 ChIP signals serve as surrogates for Pds5 binding to cohesin, as the Pds5 ChIP signal relies on cohesin binding to CARs (31). We observed that the Pds5 signal at CARs in the K112R strain was indistinguishable from that of wild type and was reduced by 2-fold at these sequences in K113R cells (Fig. S2C). These results indicated that 50% binding of Pds5 to chromatin-bound cohesin was sufficient to generate the wild-type length and position of loops in K113R cells. The Pds5 ChIP signal in Eco1-depleted (ECO1-AID) cells was also reduced by 2-fold compared to wild type (Fig. S2C). Because this level of Pds5 binding to cohesin was similar to that of K113R cohesin, the dramatic changes in the distal interactions and loss of positioned loops for unacetylated cohesin could not be attributed to the level of Pds5 binding to chromatin-bound cohesin alone.

### Evidence that acetylation of K112 reduces cohesin ATPase activity *in vitro* and *in vivo*

Several studies strongly suggested that K113 acetylation reduced Scc2/Scc4 stimulation of cohesin ATPase activity (21, 24, 25). Therefore, we wondered whether the different acetylation states of K112 might reduce cohesin ATPase activity. To compare the ATPase levels of different acetylation states, we needed homogeneous populations of cohesin acetylated at either K112, K113, or both. These purifications were not possible because Eco1 acetylation of K112 and K113 had not been reconstituted in vitro. In vivo, Eco1 acetylation of these residues was only partial in wild-type cells; therefore, cohesin purified from the K113R or K112R strains would only be partially acetylated at K112 or K113, respectively.

To overcome these hurdles, we expressed wild-type or mutant cohesins in eco1Δ wpl1Δ cells that lacked acetyltransferase activity but were viable because of the absence of Wpl1 activity (15). In eco1Δ wpl1Δ cells, the smc3-K112Q mutation (hereafter K112Q) yielded cohesin complexes containing only the K112 acetyl-mimic with no acetylation at K113. Similarly, expression of smc3-K113Q or the double mutant smc3-K112Q-K113Q in the same background generated cohesin complexes that mimicked acetylation at K113 alone (hereafter K113Q) or at both lysines (hereafter K112Q-K113Q), respectively.

We observed that Scc2/Scc4 stimulated cohesin ATPase 4.5-fold, and this stimulation was abolished by K113Q, as previously observed (21, 25, 32). The loader stimulated ATPase only 2-fold for cohesin with K112Q (Fig. 2A). These results suggested that the loader induction of cohesin ATPase was completely blocked by K113 acetylation and partially blocked by K112 acetylation.

**Fig. 2.**
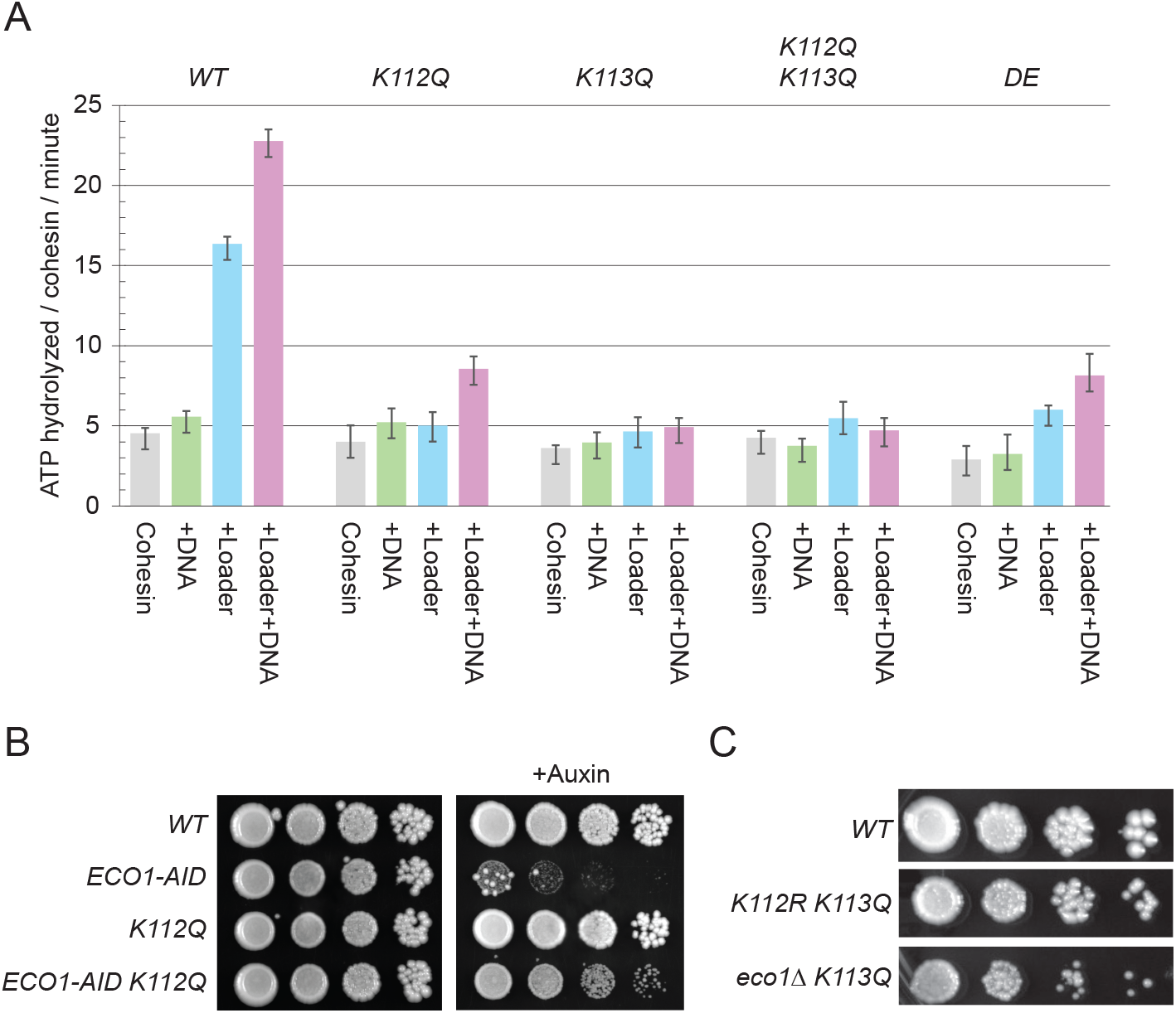
ATPase activity in wild-type and cohesin acetylation mutants. **(A)** Cohesin mutants differentially affect ATP hydrolysis rates. Wild-type and mutant cohesin complexes were purified from the acetyltransferase-defective eco1Δ wpl1Δ strains to avoid cohesin acetylation (see Table 1), and equal amounts were assessed for ATPase activities in the presence of DNA, with and without Scc2p/Scc4p (Loader). We tested ATP hydrolysis per minute for wild type complex (*WT*), single acetylation-mimic mutants smc3-K112Q (*K112Q*) and smc3-K113Q (*K113Q*), double acetylation-mimic complex (*K112Q K113Q)*, and a previously characterized mutant cohesin with a low ATPase activity, smc1-D1164E (*DE*) (strains genotypes in Table 1). **(B)** Cohesin smc3-K112Q acetyl-mimic mutant rescues viability in the absence of ECO1. Saturated cultures of cells with wild-type cohesin (*WT*), cells depleted for Eco1 acetyltransferase (*ECO1-AID)*, cells with acetyl-mimic smc3-K112Q mutant (*K112Q*), or cells depleted for Eco1 acetyltransferase and with acetyl-mimic smc3-K112Q mutant (*ECO1-AID K112Q)* (see Table 1), were spotted on rich media with or without auxin. **(C)** Cohesin smc3-K113Q lethality is rescued by blocking smc3-K112 acetylation. Saturated cultures of cells with wild-type cohesin (*WT*), cells with Smc3 acetyl-null mutant *K112R and* acetyl-mimic smc3-K113Q (*K112R K113Q)*, or cells depleted for Eco1 acetyltransferase and with acetyl-mimic smc3-K113Q mutant (*ECO1-AID K113Q)* (see Table 1), were spotted on rich media.

The reduced ATPase level of the cohesin with K112Q was similar to cohesin with the ATPase active site mutation, smc1-D1164E (henceforth referred to as DE) (Fig. 2A) (21, 25). If the K112Q mutation impacted ATPase activity in vivo as it did in vitro, the phenotypes of the K112Q mutant should be similar to the DE mutant. Like the DE mutant, the K112Q mutant had the rare ability to suppress the inviability of Eco1 depletion (Fig. 2B) (21). In addition, like the DE mutant, the K112Q mutant exhibited near wild-type levels of cohesion, robust DNA repair in response to camptothecin, and resistance to the mitotic stress of the microtubule inhibitor benomyl (Fig. S3A and B) (21, 25). These phenotypic similarities between K112Q and DE strains suggested that K112 acetylation partially represses the cohesin ATPase activity in vivo as well as in vitro.

As shown in a previous study (33), expression of the acetyl-mimic K113Q cohesin mutant alone was lethal, but viability was restored when the acetyl-null K112R mutation was added, generating the K112R K113Q double mutant (Fig. 2C). This suppression suggested that blocking acetylation at K112 counteracted the negative effect of the K113Q mutation, which mimicked acetylation. If suppression depended solely on blocking K112 acetylation, then deleting ECO1, the acetyltransferase for K112, should also restore viability to the K113Q strain, which was indeed the case (Fig. 2C). One model proposed that in the K113Q mutant, additional acetylation at K112 further reduced cohesin ATPase activity to lethal levels, and that blocking this acetylation with K112R or ECO1 deletion prevented further inhibition and rescued growth. However, this explanation is unlikely because cohesin complexes with K112Q-K113Q double mutant and K113Q alone showed the same ATPase activity (Fig. 2A). Thus, the rescue of the K113Q mutant by K112R indicates that K112 acetylation contributes to cohesin regulation through a mechanism beyond simply modulating ATPase activity.

### Impact of altering cohesin ATPase activity on chromosome structure

We performed Micro-C XL on a panel of mutants that alter cohesin ATPase activity to assess its impact on chromosome structure. This panel included cohesin with increased ATPase activity (25). The smc1-T1117I mutation (henceforth referred to as TI) had enhanced loader-induced ATPase activity. Cohesin harboring smc3-K113Q and smc1-T1117I mutations (henceforth referred to as K113Q TI) had constitutive high levels of ATPase even in the absence of the loader or DNA. The derivative slopes, decay curves, contact maps, and pileups revealed that the length distributions and positioning of loops for these strains were similar but not identical to wild type (Fig. 3 and Fig. S4). These mutants had a reduced proportion of the small loops (2–10 kb) in the decay curves (Fig. 3A and Fig. S4A), likely corresponding to the reduced random loops (the off-diagonal “fuzz”) observed in the contact maps (Fig. 3B and Fig. S4B). In contrast, the intensity of CAR-associated positioned loops increased relative to wild type (Fig. 3C and Fig. S4C). These results suggested that increased ATPase activity led to the conversion of random loops to CAR-associated positioned interactions.

**Fig. 3.**
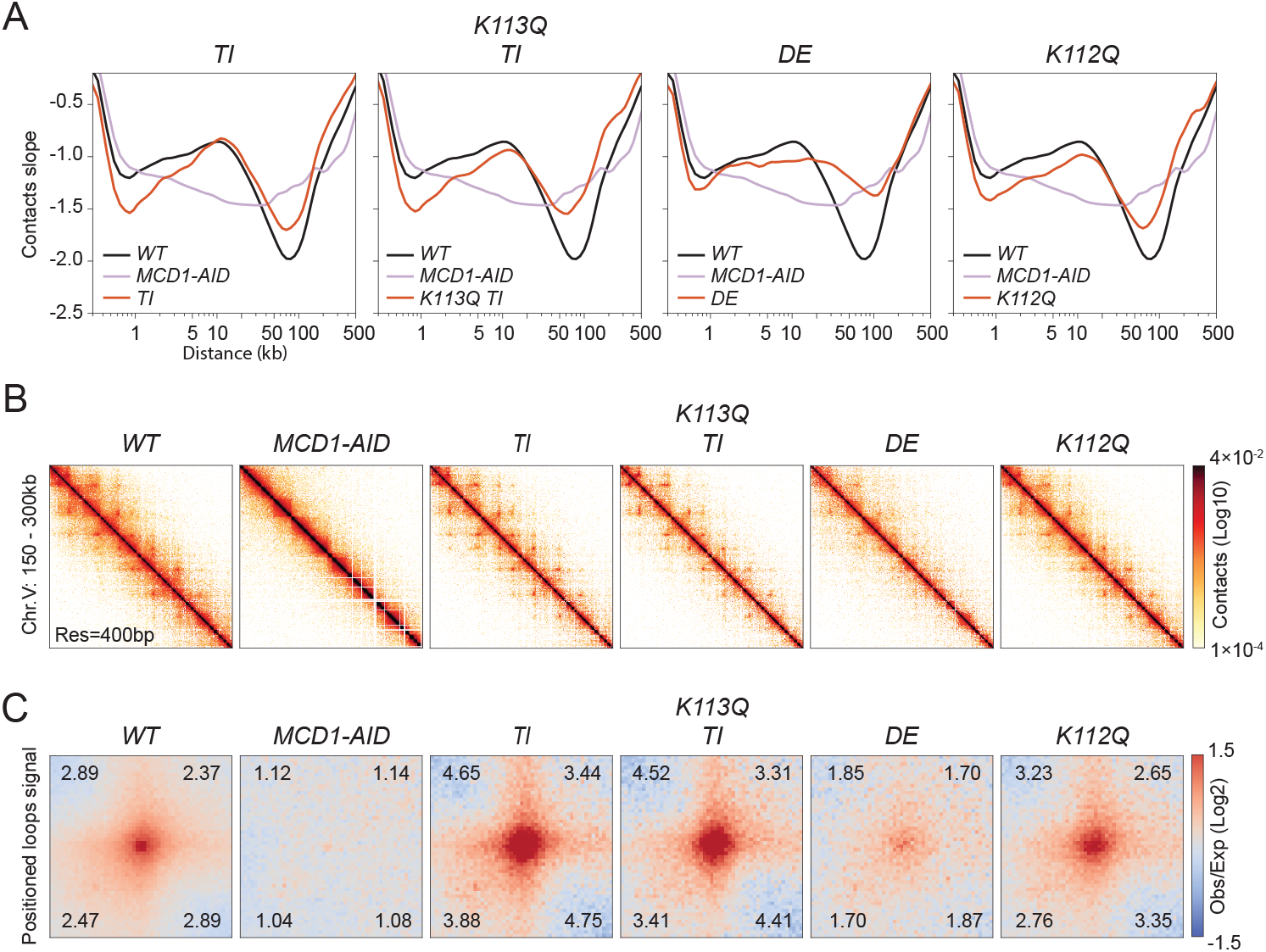
Chromosome structure in wild-type and cohesin ATPase mutants. **(A)** Chromosome contacts in cohesin ATPase mutants. Micro-C XL analysis of chromosome interactions in mitotically arrested wild-type cells (*WT*), mutants with elevated ATPase smc1-T1117I (*TI*) and smc3-K113Q combined with smc1-T1117I (*K113Q TI)*, and mutants with reduced ATPase smc1-D1164E (*DE*) and smc3-K112Q (*K112Q*) (strains genotypes in Table 1). Derivative slopes of the contact frequency curves (shown in Fig. S4A) plotted against genomic distance are depicted in black for *WT*, mauve for *MCD1-AID*, and red for ATPase mutants. **(B)** Contact maps in cohesin ATPase mutants. Micro-C XL contact maps at 400bp resolution over an arm region at chromosome V 150-300kb for *WT, MCD1-AID*, high ATPase *TI* and *K113Q TI*, and low ATPase *DE* and *K112Q* strains listed in A. **(C)** Genome-wide signal for positioned loops at CARs in cohesin ATPase mutants. Piled-up heatmap of the ±5kb regions centered at CARs for *WT, MCD1-AID*, high ATPase *TI* and *K113Q TI*, and low ATPase *DE* and *K112Q* strains listed in A. Numbers in the corners represent the fold-change of the signal enrichment of the center pixel over the indicated corner pixels.

This panel also included cohesin with decreased ATPase (DE and K112Q mutations). The K112Q decay curve revealed a length distribution of distal interactions similar to wild type, but these interactions were observed at a lower frequency (Fig. 3A and Fig. S4A). The CAR-associated, positioned loop interactions were indistinguishable from the wild type (Fig. 3B and C; Fig. S4B and C). These results suggested that cohesin with significantly lower induced ATPase was capable of generating a near-normal chromosome structure.

In contrast, the chromosome structure in the DE mutant was dramatically different from that of the wild type or the K112Q mutant. The DE mutant showed a reduction in interactions around the 10 kb range and a corresponding increase in distal interactions spanning 50–150 kb (Fig. 3A and Fig. S4A). The contact maps and pile-ups revealed that positioned loops decreased dramatically (Fig. 3B and C; Fig. S4B and C). These results suggested that in the DE mutant, looping failed to stop at CAR sites, generating much longer loops with loop anchors at random genomic positions. The loss of positioned loops cannot be explained by the reduced ATPase activity of DE cohesin because the K112Q cohesin had equally reduced ATPase but formed positioned loops like wild type (Fig. 2A and Fig. 3). Furthermore, the fact that DE cohesin could form larger loops than wild type despite having lower ATPase activity suggested loop size in wild-type cells was not limited by cohesin’s ATPase activity.

To further investigate the mechanism of DE’s longer loops, we leveraged the fact that longer distal interactions had also been observed previously upon depletion of Wpl1 or Eco1 (11, 22). Several observations suggest that these longer loops arise through distinct mechanisms in each mutant. First, analysis of contact decay curves showed that the double mutant for Eco1 and Wpl1 (eco1Δ wpl1Δ) displayed even longer-range interactions than cells lacking only Eco1 (ECO1-AID) or Wpl1 (wpl1Δ) (Fig. 4A and Fig. S5A). Second, Eco1 depletion, unlike Wpl1 depletion, caused the loss of positioned loops (Fig. 4A and Fig. S5B–D) (11, 22). Interestingly, DE mutants phenocopied Eco1-depleted cells: both lacked positioned loops (Fig. S5B–D) and exhibited highly similar contact decay curves and derivative slopes (Fig. 4A and Fig. S5A). In contrast, DE cells did not resemble the double eco1Δ wpl1Δ mutants, which showed even more extensive looping. Together, these findings suggest an epistatic relationship between the DE mutation and Eco1 depletion. Specifically, DE and Eco1 depletion act in a shared pathway that constrains loop length and supports loop positioning, distinct from the pathway affected by Wpl1.

**Fig. 4.**
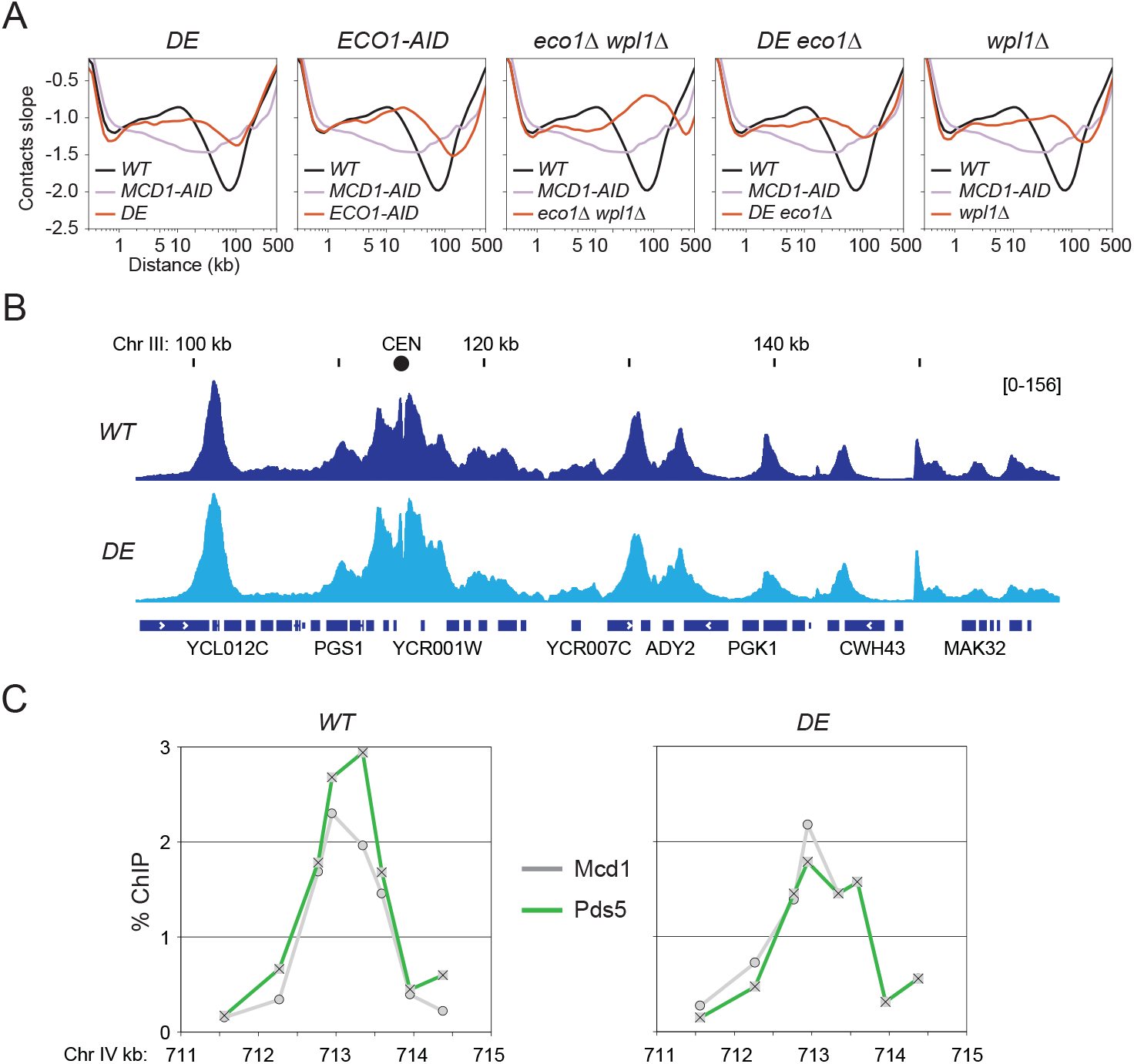
Epistasis analysis of mutants with expanded loops. **(A)** Chromosome contacts in cohesin mutants with expanded loops. Micro-C XL analysis of chromosome interactions in mitotically arrested wild-type cells (*WT*), cells depleted for cohesin (*MCD1-AID*), cells with smc1-D1164E (*DE*), cells depleted of Eco1 acetyltransferase (*ECO1-AID*), cells deleted for Eco1 and Wpl1 (*eco1*Δ *wpl1*Δ), cells with smc1-D1164E combined with Eco1 deletion (*DE eco1*Δ), cells deleted for Wpl1 (*wpl1*Δ) (strains genotypes in Table 1). Derivative slopes of the contact frequency curves (shown in Fig. S5A) plotted against genomic distance are depicted in black for *WT*, mauve for *MCD1-AID*, and red for the indicated mutants. **(B)** Cohesin smc1-D1164E mutant binds DNA with a wild-type profile. ChIP-seq profile of mitotically arrested *WT* and *DE* cells. **(C)** Pds5 binds to the cohesin smc1-D1164E mutant on chromatin. ChIP qPCR profile of mitotically arrested *WT* and *DE* cells for Mcd1 (grey) and Pds5 (green) on a representative CAR.

Cohesin acetylation is known to stabilize its binding to DNA (20), and such stable binding could act as a barrier to translocating cohesin, thereby constraining loop size and promoting the formation of positioned loops. This model could explain why Eco1-depleted cells, which lack cohesin acetylation, exhibit both enlarged loops and a loss of positioned loops. Based on this reasoning, the DE mutant, which also displays long loops and reduced positioned loops, might similarly destabilize cohesin– DNA interactions. However, several lines of evidence argue against this possibility. First, ChIP-seq analysis revealed that the genome-wide pattern and intensity of cohesin binding in DE cells are comparable to wild type (Fig. 4B) (21). Second, we directly tested the stability of cohesin binding at CARs. Cohesin was allowed to bind DNA during S phase, cells were then arrested in mitosis, and the Scc2 loader was inactivated to prevent new cohesin loading. We then monitored the levels of CAR-bound cohesin prior to and after loader inactivation (Fig. S6A). In DE cells, cohesin levels remained unchanged at an arm CAR and decreased by ~50% at centromeres after one hour—a stability profile indistinguishable from wild type (Fig. S6B). Moreover, cohesin binding stability correlates with cohesion maintenance. The DE mutant supports near-wild-type levels of cohesion, consistent with previous reports (17, 18, 21, 25). Finally, positioned loops were not observed in eco1Δ wpl1Δ cells, where cohesin binding to chromosomes was stabilized by Wpl1 inactivation (20, 22, 23). Together, these findings argue that destabilized cohesin binding is not the shared cause of long, random loops in DE or Eco1-depleted cells.

Recent studies have proposed that Eco1-mediated acetylation constrains the length of cohesin-dependent loops by promoting the recruitment of Pds5 to chromatin-bound cohesin (22, 23). Based on this model, we hypothesized that the DE mutant could impact this mechanism by either preventing Eco1 from acetylating cohesin or altering cohesin such that acetylation can no longer enhance Pds5 binding. We tested the first possibility by looking at K113 acetylation levels using a K113ac-specific polyclonal antibody. We found that K113 acetylation was maintained in the DE mutant at levels comparable to wild type (Fig. S6C), ruling out a defect in acetylation. We tested the second possibility by examining Pds5 binding to CARs in the DE mutant by ChIP (Fig. 4C). No significant difference in the Pds5 ChIP signal was observed between wild type and DE, suggesting that recruitment of Pds5 to cohesin was intact in the DE mutant. These results indicate that the defect in the DE mutant lies downstream of Pds5 recruitment. We propose that the DE mutation alters cohesin in a way that renders it unresponsive to Pds5-dependent modulation—highlighting a previously unrecognized step in the pathway that governs the formation of positioned loops.

## Discussion

By studying a panel of cohesin mutants in budding yeast, we provide important new insights into the functions and interplay of cohesin ATPase and acetylation in tethering and looping.

### The role of Smc3 K113 acetylation in tethering

Our analysis of the K113R acetylation-null mutant supports a model in which sister chromatid cohesion is established through the sequential capture of sisters (21, 34, 35). We show that the K113R mutant exhibits a near wild-type distribution of cohesin binding across the genome, as well as normal size and positioning of cohesin-dependent loops. Similar observations were reported for a cohesin hinge mutant in mammalian cells (36). Yet these mutants fail to generate cohesion. Previously, it was hypothesized that the failure of such mutants to establish cohesion might stem from unstable cohesin–DNA binding, thereby impairing cohesion maintenance (15, 37). However, subsequent studies in yeast and human cells showed that cohesin in K112R-K113R double mutants bound stably to DNA, yet still failed to generate cohesion (37, 38). Thus, the failure of the K113R mutant to support cohesion is not due to an inability to stably bind one sister chromatid, but rather a failure to capture or stably tether the second sister. These findings indicate that K113 acetylation is essential for the successful engagement of the second sister chromatid, and thereby, for the establishment of cohesion.

### The role of cohesin tethering in looping

A second insight from our mutant panel study addresses the mechanism underlying chromatin loop formation in budding yeast. A recent study proposed that cohesin-dependent distal interactions may arise through a mechanism distinct from loop extrusion. In this alternative model, termed loop capture, cohesin first topologically loads onto DNA and then sequentially entraps a second DNA segment, forming a loop. Instead of active extrusion, loop formation happens passively, with transcription helping move cohesin and bring distant sites together (39). This model makes several strong predictions. First, entrapment-competent cohesin that generates cohesion and binds to wild-type sites should produce loops with wild-type length and positioning. However, we showed that the DE mutant forms longer distal interactions with a near-complete loss of positioned loops, while binding to chromatin like wild type and exhibiting robust entrapment/cohesion activity (21, 25). This suggests that topological entrapment alone is not sufficient to constrain loop size or positioning in vivo.

An even stronger prediction of the loop capture model is that cohesin mutants defective in entrapping the second DNA sequence should fail both cohesion and loop formation. However, the K113R mutant, which binds chromatin as wild type and is cohesion-defective, still generates loops with length and positioning similar to wild type. A similar result was reported for a cohesin hinge mutant (35). Taken together, these findings indicate that loop formation does not require the tethering activity that mediates cohesion, and instead support a model in which loops are generated through loop extrusion.

### The role of Smc3 K112 acetylation in ATPase regulation and cohesin function

Previous studies failed to demonstrate a function for the conserved acetylation of Smc3 at K112. Our studies of the K112 and K113 acetyl mimics suggest that K112 acetylation causes partial inhibition of Scc2-induced cohesin ATPase activity in vitro and in vivo (this study), in contrast to its complete inhibition by K113 acetylation (this study; 24, 25). These results indicate that differential acetylation of K112 and K113 enables cells to generate two distinct Scc2-responsive states of cohesin ATPase: non-inducible (K113 acetylation) or partially inducible (K112 acetylation).

The biological significance of the partially inducible ATPase state conferred by K112 acetylation remains unclear. We found that the K112R acetyl-null mutant behaves like wild type across a range of molecular and cellular assays, including cohesin binding to DNA, loop positioning and length, cohesion, and resistance to camptothecin and benomyl, which serve as proxies for DNA repair and chromosome segregation, respectively (this study). One approach to uncover the biological function of K112 acetylation is to identify mutations that generate synthetic phenotypes in combination with the acetyl-null K112R mutation. This approach may reveal yet another cohesin function and provide a paradigm for studying the abundant but poorly understood posttranslational modifications of ABC ATPase proteins.

### The role of Smc3 K112 and K113 acetylation in looping

Our analysis of cohesin acetylation mutants revealed unexpected complexity in the mechanisms that govern the formation of positioned chromatin loops. We demonstrated that positioned loops formed in the K112R and K113R single mutants (this study), in contrast to their loss in the K112R-K113R double mutant (22). This suggests that acetylation of either K112 or K113 is necessary to support loop positioning, indicating a degree of functional overlap between the two modifications. One possible explanation is that acetylation at either site reduces cohesin ATPase activity (this study), which could stop loop extrusion, thus forming positioned loops. However, our data show that the DE mutant, which also exhibits low ATPase activity comparable to K112-acetylated cohesin (K112Q), is significantly impaired in forming positioned loops. This finding suggests that while reduced ATPase activity may be necessary, it is not sufficient for the establishment of positioned loops. Therefore, K112 and K113 acetylation must influence loop positioning through mechanisms beyond ATPase inhibition alone.

A compelling hypothesis is that K112 or K113 acetylation promotes loop positioning by enhancing cohesin’s interaction with Pds5, a factor known to bind acetylated cohesin (40), and shown to be required for positioned loop formation (11, 22, 41). However, in our study, DE cohesin was acetylated, bound Pds5 robustly, yet failed to generate positioned loops, indicating that Pds5 binding alone is also not sufficient. We propose that Pds5 must influence cohesin function or conformation. The DE mutation, which was originally selected to bypass the need for acetylation, may confer a cohesin conformation that is refractory to modulation by Pds5, thereby preventing proper loop positioning. These findings point to a more nuanced regulatory role for acetylation—beyond ATPase inhibition or cofactor recruitment—in shaping cohesin’s loop-forming capacity.

### The role of cohesin ATPase in looping

Our cohesin mutant panel provided important insights into how ATPase activity influences chromosome structure. Cohesin complexes with elevated ATPase activity, such as those carrying the smc1-T1117I (TI) or smc3-K113Q with smc1-T1117I (K113Q TI) mutations, exhibited fewer randomly occurring loops and more positioned loops compared to wild type. One explanation for this pattern is that loop extrusion proceeds dynamically until cohesin encounters a boundary element that halts translocation and stabilizes a positioned loop, while random loops represent transient intermediates formed during this process. In this model, cohesin with higher ATPase activity may translocate more rapidly, spending less time in the intermediate positions. Alternatively, random loops may result from abortive looping attempts, and higher ATPase activity could enhance processivity, enabling cohesin to reach stop signals and establish stable loops more reliably. Finally, high ATPase activity may stabilize cohesin at boundary elements, thereby allowing for the accumulation of more positioned loops. These hypotheses can be tested in future in vitro loop extrusion assays.

Interestingly, we found that mutations that reduce cohesin ATPase activity did not necessarily impair the formation of chromatin loops. For example, the K112Q mutant produced distal interactions similar in length to wild type, while the DE mutant generated longer-range interactions. This raises the possibility that ATP hydrolysis may not be the rate-limiting step in the loop extrusion process, implying that ATP hydrolysis may not control how quickly or how far cohesin extrudes loops. However, there is an important caveat to this interpretation. Our experiments used Micro-C XL assays performed after mitotic arrest, meaning that cells were held in a paused state for an extended period. This pause could have allowed even slowly functioning cohesin complexes more time to extrude loops, potentially masking defects in loop extrusion speed. To determine whether ATP hydrolysis truly influences loop length and speed, it will be important to perform additional analyses—including time-resolved studies both in vivo and in vitro—to directly measure cohesin behavior under dynamic conditions.

## Acknowledgments

This work was funded by a National Institutes of Health grant, 1R35 GM-118189-06 (to D.K.) and NIH grant R35GM133678 (to W.M.). The work of L.C. was funded by the Austrian Academy of Sciences (ÖAW). We thank Jan-Michael Peters at IMP Vienna for supporting L.C.

## Author contributions

L.C. designed and performed Micro-C experiments.

T.Y. and Y.M. performed the Micro-C computational analyses. K.B. and J.L. performed the ChIP-seq and ChIP-qPCR. S.X. designed and performed the ATPase activities. W.M. provided support for the computational analysis. D.K., and L.C. wrote the manuscript with editing comments provided by K.B., S.X., and W.M.

## Methods

### Yeast strains, media, and reagents

Yeast strains used in this study are A364A background unless otherwise specified. Genotypes are listed in Supplemental Table 1. YPD media was prepared as previously described (Guacci et al., 1997). Plates containing benomyl or camptothecin (Sigma catalog # C9911) were used to assess drug sensitivity, as previously described (Guacci and Koshland 2012). Auxin (3-indoleacetic acid) (Sigma-Aldrich, St. Louis, MO) was prepared as a 1 M stock solution in DMSO and then added to the liquid media or plates at a final concentration of 500 μM or 750 μM, respectively.

Cohesin Purification Media: Low Biotin Synthetic Complete (LBSC) Media contained 1.56 g/L BSM Powder (Sunrise Science Products Cat#1387), 1.71 g/L YNB – Biotin powder (Sunrise Science Products Cat#1523), 38 mM ammonium sulfate (5 g/L), 1 nM D-biotin (Invitrogen #B20656), and 2% raffinose.

Cohesin Loader Purification Media: Low Biotin URA-Dropout Media contained 0.8g/L CSM-Ura (Sunrise Science Products), 1.71g/L YNB –Biotin powder (Sunrise Science Products Cat#1523), 38 mM ammonium sulfate (5g/L), 1 nM biotin, and 2% raffinose.

CRISPR guide plasmids and PCR-generated repair templates were made and used to insert mutations into yeast as previously described (Saxton and Rine 2019).

Preparation of mitotically arrested cells for Micro-C XL, ChIP, and cohesion assays. Asynchronous mid-log cultures were arrested in G1 by the addition of alpha factor as previously described (Guacci et al. 2019). When required, auxin was added (500 μM final) to G1-arrested cells, incubated for 30 minutes while arrested in G1. G1-arrested cells were released from G1 into either YPD or YEPRG containing nocodazole and Pronase E as previously described (Guacci et al. 2019), then incubated at 30°C for 2.5h for YPD or 4h for YEPRG to arrest in mid-M phase. When required, auxin was added (500 μM final) in all wash media and in resuspension media to ensure AID-tagged protein depletion.

### Protein extracts and western blotting

Protein extracts and western blots for detecting Mcd1, Scc2-V5, and Smc3-K113 acetylation were performed as described previously (Eng et al. 2014; Robison et al. 2018).

### Cohesion Assay

Cohesion was monitored at LYS4 using the LacO-LacI system as previously described (Guacci and Koshland 2012). Mid-M phase cells were fixed, and the number of GFP signals in each cell was scored. Cells with 2 GFP spots have defective cohesion.

### Chromatin Immunoprecipitation (ChIP) and Micro-C XL

ChIP-qPCR, ChIP-seq, and Micro-C XL were prepared and analyzed as described previously (Eng et al. 2014; Costantino et al. 2020)

### Purification of cohesins and ATPase assay

Purifications of loader (SX305) and cohesin from wild type (KB58A) and mutants (SX364, KB140A, SX365, and SX316) were performed as described previously (Boardman et al. 2023). ATPase activity of cohesin was measured using EnzChek Phosphate Assay Kit with purified recombinant proteins depleted of free phosphate using Inorganic Phosphate Binding Resin (Abcam: ab270547). Reactions were assembled with 10nM cohesin, 15nM Scc3, alone or with 65nM Scc2/4, 0.1mg/ml BSA, and 450nM 60-mer dsDNA in ATPase reaction buffer (25mM HEPES pH7.5, 20% glycerol, 50 mM NaCl, 1 mM MgCl2); reactions were initiated with the addition of ATP to a final concentration of 1mM. Spectrophotometric measurements at 360 nm were taken every 1 min for 2h at room temperature. ATPase activities were calculated by linear regression of the raw data using GraphPad Prism software.

### Processing Micro-C XL reads to contact matrices

Each Micro-C XL fastq raw file was processed into contact read pairs using HiC-Pro version 3.1.0 pipeline (https://github.com/nservant/HiC-Pro, Servant et al., 2015). Reads were aligned to the yeast sacCer3 genome using Bowtie2 version 2.4.4 (Langmead and Salzberg, 2012) with ‘--very-sensitive-local’ option. Singleton reads, multi-mapped reads, and PCR duplicate read pairs were discarded. Aligned read pairs with genomic distances shorter than 200 bp were also filtered out. The output valid contact pairs were binned at multiple resolutions and converted to contact matrices in both .cool format using cooler (https://github.com/open2c/cooler, Abdennur and Mirny, 2020) and .hic format using Juicer (https://github.com/aidenlab/juicer, Durand et al., 2016) for downstream analyses. Contact frequency matrices in .cool format were normalized using IC (iterative correction) (Imakaev et al., 2012) via the cooler balance command, and matrices .hic format were normalized by KR (Knight-Ruiz) balancing (Knight and Ruiz, 2013) using Juicer. The MCD1-AID mutant contact matrices in both .cool and .hic format were downloaded from GEO (GSE151553, Costantino et. al, 2020). Hi-C datasets for WT (CH112, CH233, CH283) and Smc3 RR, Smc3-AID (CH212, CH287) (Bastié et al., 2022) were downloaded from the SRA database (PRJNA715343). Hi-C fastq files were processed in a similar way as Micro-C XL data, with an additional step to generate restriction fragments after DpnII + HinfI restriction enzymes digestion. Only aligned reads that could be assigned to a restriction fragment were retained. Replicates were merged and normalized into single contact matrices in both .cool and .hic format. Heatmaps of contact matrices were plotted at 400 bp resolution using matplotlib version 3.8.3 (Hunter, 2007).

### Contact frequency versus genomic distance decay curves analysis

We used intra-chromosomal ‘UNI’ orientation contact read pairs binned at 100 bp resolution to calculate the average contact frequency between pairs separated by the same genomic distance range (decay curves), using the expected module from cooltools version 0.6.1 (https://github.com/open2c/cooltools, Open2C, et al., 2024). The ‘UNI’ orientation is defined as aligned read pairs mapping to the same strand direction (either +/+ or −/−) (Hsieh et al., 2015). Filtered pairs were then converted into contact matrices in .cool format. The average contact frequencies at each genomic distance (matrix diagonals) were calculated independently for each chromosome using the diagsum_symm function from cooltools. To account for the sparsity of contacts at larger genomic distances, the data were grouped into logarithmically spaced bins (20 bins per order of magnitude) using logbin_expected function from cooltools. Genome-wide averages of log-binned average contact frequencies and decay slopes were then computed by combining chromosome-level values with combine_binned_expected function from cooltools. Decay curves, slopes, and slope differences (Δslope) were plotted using matplotlib. Micro-C XL and Hi-C decay curves were normalized at 1 kb and 3.2 kb distances, respectively.

### Chromatin loops and pileup analysis

Loops were identified using the HiCCUPS algorithm (Rao et al., 2014) implemented in Juicer. We applied the algorithm to KR-normalized contact matrices in .hic format at 500 bp resolution, and the results were filtered at 1% false discovery rate. Loops were called at two settings, then merged: peak/window width (6,12) and (8,16). Pixels within 2500 bp of each other were merged. The calling options were: hiccups -m 4096 -k KR -r 500,500 -f 0.1,0.1 -p 6,8 -i 12,16 -d 2500,2500 (Costantino et al., 2020). Loop anchors separated by distance greater than 100 kb were considered likely false positives and discarded. Genome-wide positional loop signals were quantified using aggregate peak analysis (Rao et al., 2014). Pileups were generated on IC-normalized contact matrices in .cool format at 200 bp resolution using coolpup.py version 1.1.0 (https://github.com/open2c/coolpuppy, Flyamer et al., 2020). Pileups were centered at the wild-type loop anchors with ±5 kb flanking regions. To account for distance-dependent decay effects, pileups were normalized against background contact levels using randomly shifted control regions. The calling options were: minshift=1000, maxshift=10000, flank=5000, nshifts=10. Loop enrichment was calculated as the ratio of the mean normalized center contacts (within a 5×5 window) to the mean normalized corner contacts. Pileups were plotted with matplotlib.

The same aggregate peak analysis was applied to measure target-centered loop signals, using paired MCD1 ChIP-seq peaks (CARs) instead of HiCCUPS-called loop anchors. ChIP-seq raw fastq files were downloaded from the SRA database (SRR11872088 and SRR11872089, Costantino et. al, 2020). Reads were aligned to the yeast sacCer3 genome using Bowtie2. Peaks were called with MACS2 version 2.2.7 (Zhang et al., 2008) with default settings, and replicates were combined using IDR version 2.0.4 (https://github.com/nboley/idr, Li et al., 2011) at a threshold of 0.05. MCD1 peaks were paired by genomic intervals ranging from +1 to +10, where +1 denotes the directly adjacent peaks.

---

**Fig. S1.**
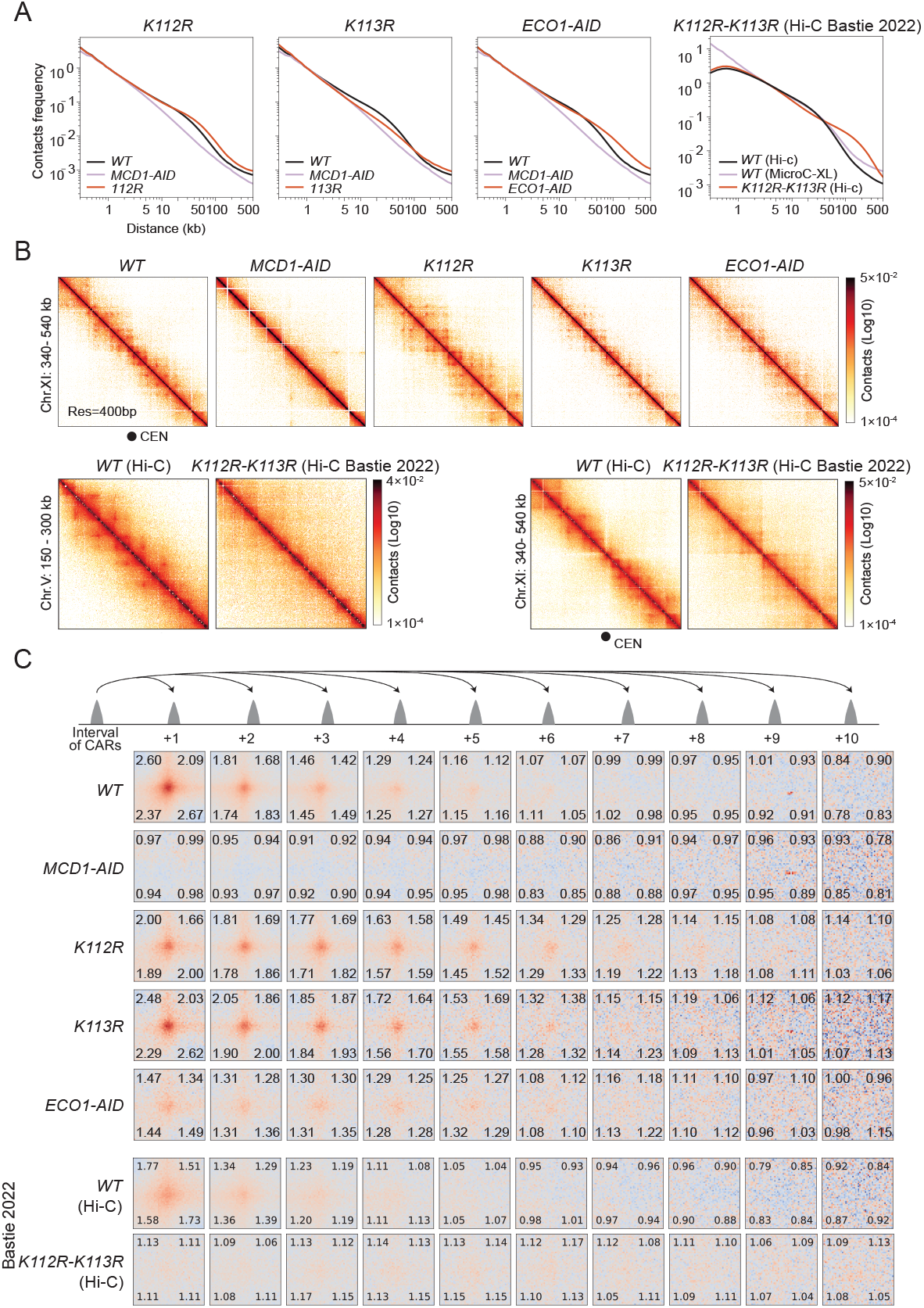
Chromosome structure in wild-type and cohesin acetylation mutants. **(A)** Chromosome contacts in cohesin acetylation mutants. Micro-C XL analysis of chromosome interactions in mitotically arrested wild-type cells (*WT*), single acetylation-deficient mutants smc3-K112R (*K112R*) and smc3-K113R (*K113R*), and cells depleted of Eco1 acetyltransferase (*ECO1-AID*) or cohesin (*MCD1-AID*) (strains genotypes in Table 1). Interactions-versus-distance decaying curve shows the normalized contact density (y-axis) against the distance between the pair of crosslinked nucleosomes from 100bp to 1Mb (x-axis). *WT* is depicted in black, *MCD1-AID* in mauve, and acetylation-mutants in red. Wild type (*WT* Hi-C) and smc3-K112R-K113R (*K112R-K113R* Hi-C) from Hi-C experiments were also plotted (Bastié et al. 2022. **(B)** Contact maps in cohesin acetylation mutants over a centromere. Micro-C XL contact maps at 400bp resolution over the centromeric region at chromosome XI 340-540kb for the *WT, MCD1-AID, K112R, K113R*, and *ECO1-AID* strains listed in A. The centromere position is depicted with a black circle. Wild type (*WT* Hi-C) and smc3-K112R-K113R (*K112R-K113R* Hi-C) from Hi-C experiments were also plotted (Bastié et al. 2022). **(C)** Genome-wide signal for positioned loops at different CAR intervals in cohesin acetylation mutants. Piled-up heatmap of the ±5kb regions centered at different intervals of CARs from +1 to +10 for the *WT, MCD1-AID, K112R, K113R*, and *ECO1-AID* strains listed in A. Numbers in the corners represent the fold-change of the signal enrichment of the center pixel over the indicated corner pixels. Wild type (*WT* Hi-C) and smc3-K112R-K113R (*K112R-K113R* Hi-C) from Hi-C experiments were also plotted (Bastié et al. 2022).

**Fig. S2.**
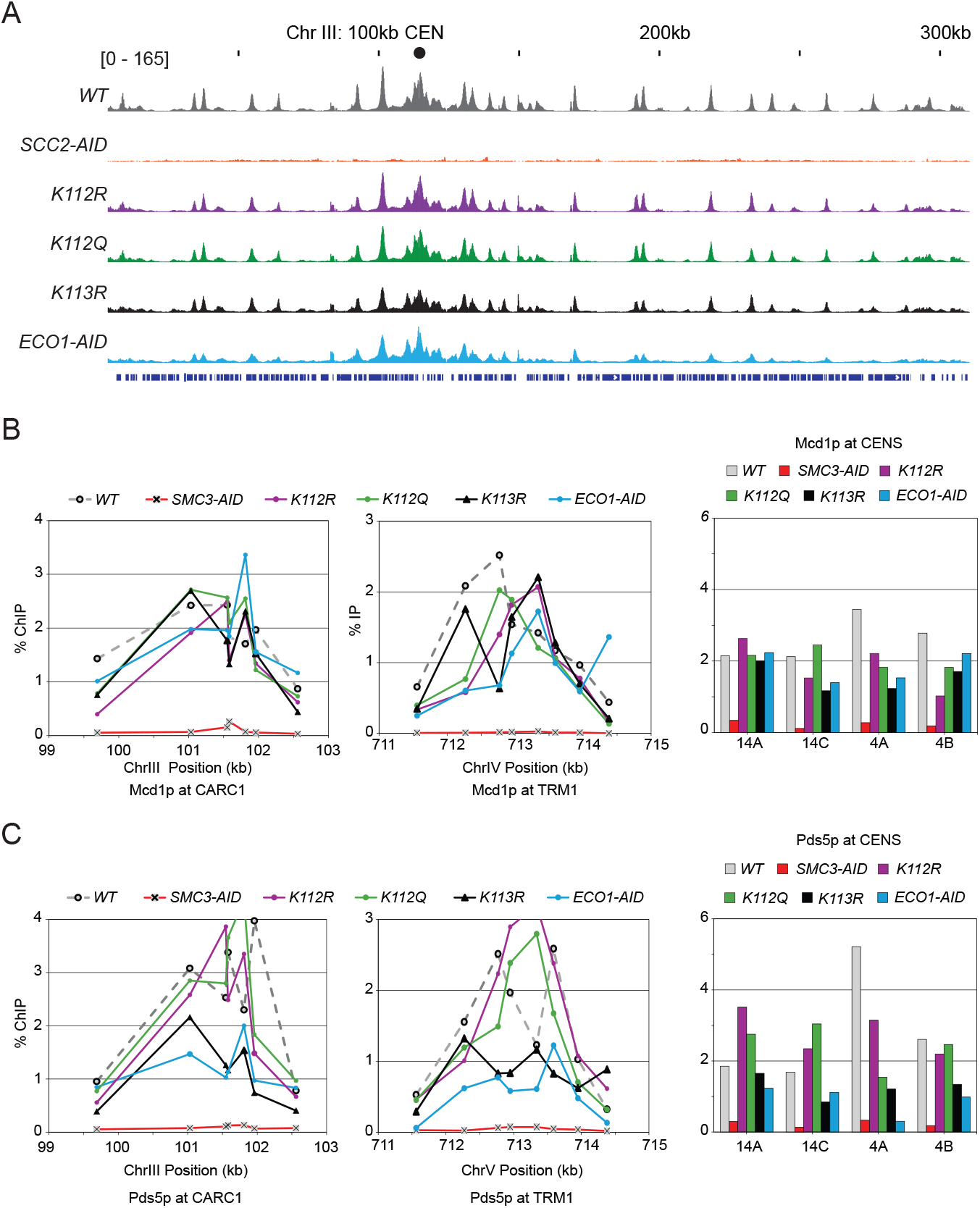
Cohesin acetylation mutants bind chromatin as wild type. **(A)** Cohesin acetylation mutants bind chromatin as wild type by ChIP-seq. ChIP-seq profile of mitotically arrested wild-type cells (*WT*), cells depleted for Scc2 loader (*SCC2-AID*), cells with acetyl-null smc3-K112R (*K112R*) mutant, cells with acetyl-mimic smc3-K112Q (*K112Q*), cells with acetyl-null smc3-K113R (*K113R*), and cells depleted of Eco1 acetyltransferase (*ECO1-AID*) (strains genotypes in Table 1). **(B)** Cohesin acetylation mutants can bind CARs and centromeres by ChIP-qPCR. ChIP qPCR of mitotically arrested *WT, SCC2-AID, K112R, K112Q, K113R*, and *ECO1-AID* strains listed in A, for Mcd1 binding at two representative CARS, CARC1 (right), TRM1 (center), and indicated centromeres (left). **(C)** Pds5 can bind to cohesin acetylation mutants at CARs and centromeres by ChIP-qPCR. ChIP qPCR of mitotically arrested *WT, SCC2-AID, K112R, K112Q, K113R*, and *ECO1-AID* strains listed in A, for Pds5 binding at two representative CARS, CARC1 (right), TRM1 (center), and indicated centromeres (left).

**Fig. S3.**
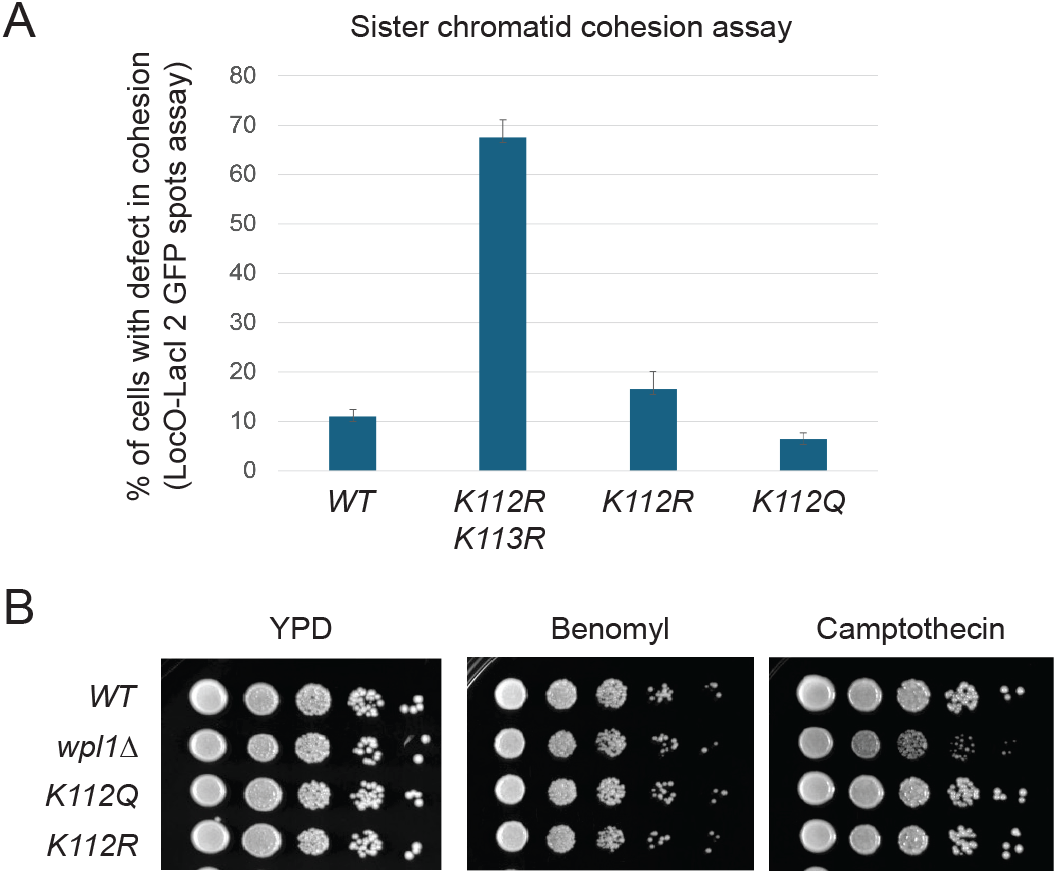
Cohesin smc3-K112Q mutant in cohesion and drug sensitivity. **(A)** Cohesin smc3-K112Q mutant presents wild-type levels of sister chromatid cohesion. Mitotically arrested cells were scored for cohesion defect (separated 2 GFP spots) for wild-type (*WT*), cells with smc3-K112R and smc3-K113R (*K112R K113R*), cells with smc3-K112R (*K112R*), and cells with smc3-K112Q (*K112Q*) (strains genotypes in Table 1). **(B)** Cohesin smc3-K112Q mutant shows robust DNA repair and resistance to mitotic stress. Saturated cultures of cells with wild-type cohesin (*WT*), cells deleted for Wpl1 (*wpl1*Δ), cells with smc3-K112Q (*K112Q*), and cells with smc3-K112R (*K112R*) (strains genotypes in Table 1) were spotted on rich media with or without benomyl or camptothecin.

**Fig. S4.**
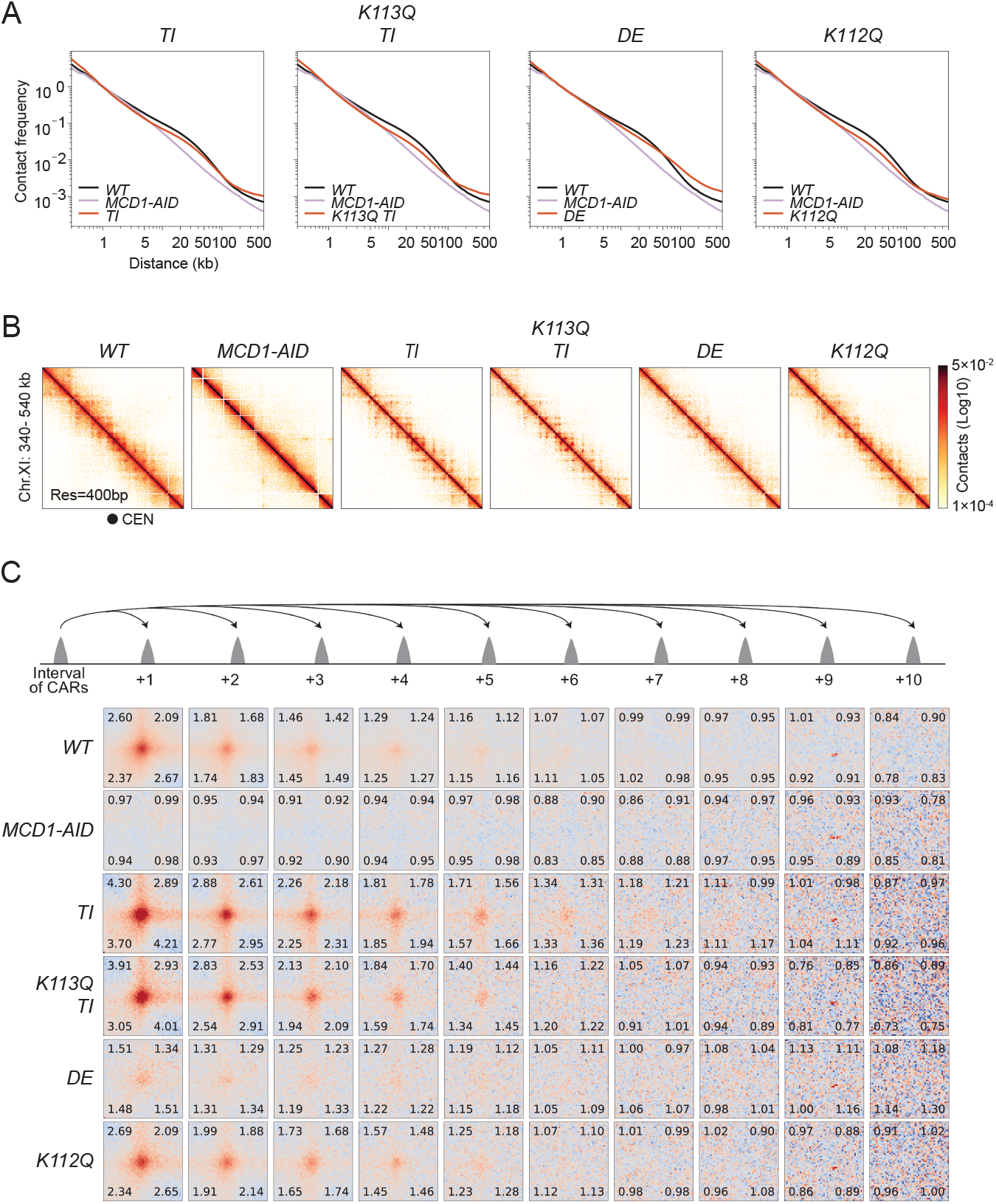
Chromosome structure in wild-type and cohesin ATPase mutants. **(A)** Chromosome contacts in cohesin ATPase mutants. Micro-C XL analysis of chromosome interactions in mitotically arrested wild-type cells (*WT*), mutants with elevated ATPase as smc1-T1117I (*TI*) and smc3-K113Q combined with smc1-T1117I (*K113Q TI)*, and mutants with reduced ATPase as smc1-D1164E (*DE*) and smc3-K112Q (*K112Q*) (strains genotypes in Table 1). Interactions-versus-distance decaying curve shows the normalized contact density (y-axis) against the distance between the pair of crosslinked nucleosomes from 100bp to 1Mb (x-axis). *WT* is depicted in black, *MCD1-AID* in mauve, and ATPase mutants in red. **(B)** Contact maps in cohesin ATPase mutants over a centromere. Micro-C XL contact maps at 400bp resolution over the centromeric region at chromosome XI 340-540kb for the *WT, MCD1-AID*, high ATPase *TI* and *K113Q TI*, low ATPase *DE* and *K112Q* strains listed in A. The centromere position is depicted with a black circle. **(C)** Genome-wide signal for positioned loops at different CAR intervals in cohesin ATPase mutants. Piled-up heatmap of the ±5kb regions centered at different intervals of CARs from +1 to +10 for the *WT, MCD1-AID*, high ATPase *TI* and *K113Q TI*, low ATPase *DE* and *K112Q* strains listed in A. Numbers in the corners represent the fold-change of the signal enrichment of the center pixel over the indicated corner pixels.

**Fig. S5.**
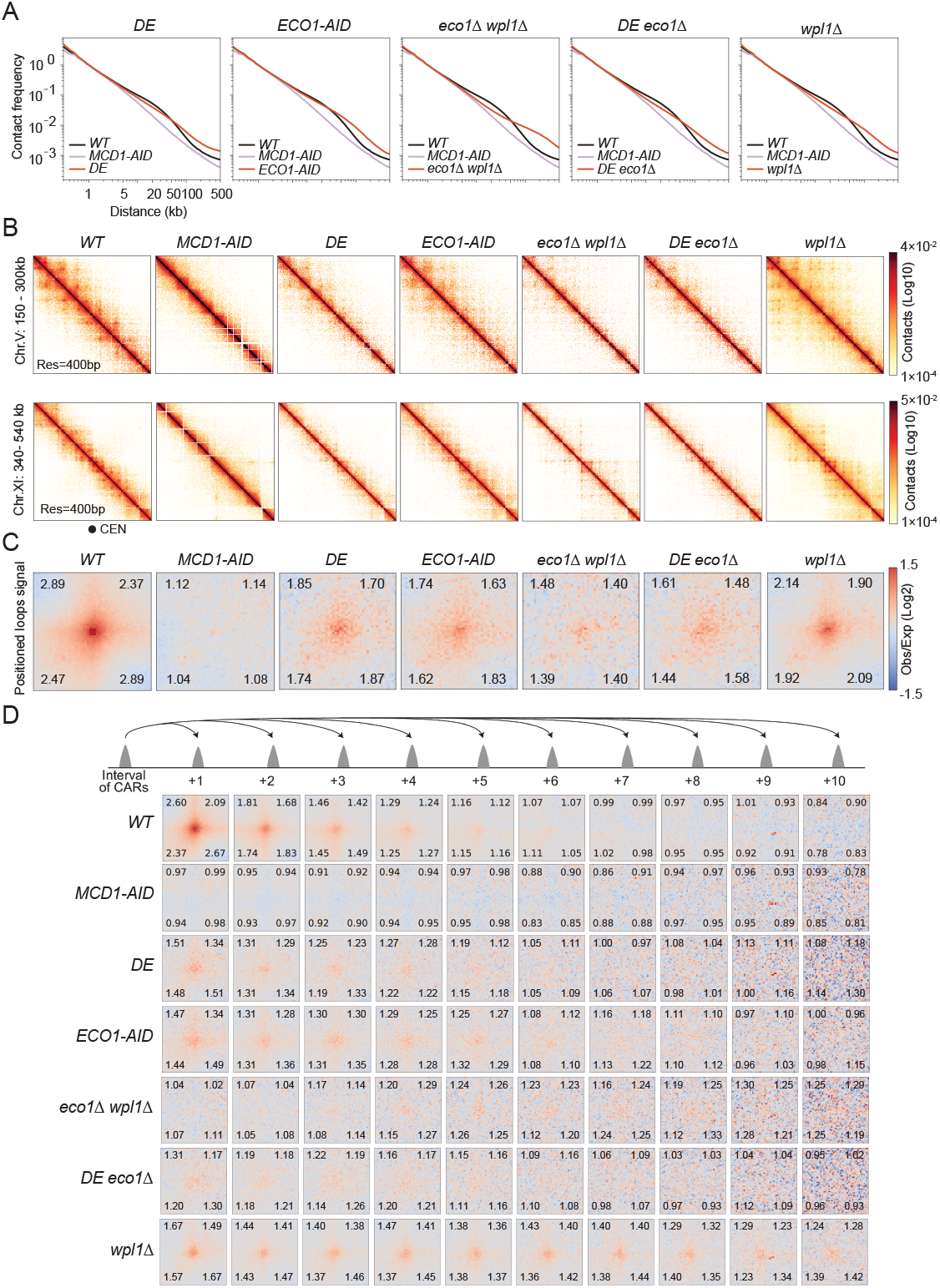
Chromosome structure of mutants with expanded loops. **(A)** Chromosome contacts in mutants with expanded loops. MiMicro-C XL analysis of chromosome interactions in mitotically arrested wild-type cells (*WT*), cells depleted for cohesin (*MCD1-AID*), cells with smc1-D1164E (*DE*), cells depleted of Eco1 acetyltransferase (*ECO1-AID*), cells deleted for Eco1 and Wpl1 (*eco1*Δ *wpl1*Δ), cells with smc1-D1164E combined with Eco1 deletion (*DE eco1*Δ), cells deleted for Wpl1 (*wpl1*Δ) (strains genotypes in Table 1). Interactions-versus-distance decaying curve shows the normalized contact density (y-axis) against the distance between the pair of crosslinked nucleosomes from 100bp to 1Mb (x-axis). *WT* is depicted in black, *MCD1-AID* in mauve, and mutants in red. **(B)** Contact maps in mutants with expanded loops over a chromosome arm and a centromere. Micro-C XL contact maps at 400bp resolution over an arm region at chromosome V 150-300kb (top), and over the centromeric region at chromosome XI 340-540kb (bottom) for the *WT, MCD1-AID, DE, ECO1-AID, eco1*Δ *wpl1*Δ, *DE eco1*Δ, *wpl1*Δ strains listed in A. The centromere position is depicted with a black circle. **(C)** Genome-wide signal for positioned loops at CARs in mutants with expanded loops. Piled-up heatmap of the ±5kb regions centered at CARs for *WT, MCD1-AID, DE, ECO1-AID, eco1*Δ *wpl1*Δ, *DE eco1*Δ, *wpl1*Δ strains listed in A. Numbers in the corners represent the fold-change of the signal enrichment of the center pixel over the indicated corner pixels. **(D)** Genome-wide signal for positioned loops at different CAR intervals in mutants with expanded loops. Piled-up heatmap of the ±5kb regions centered at different intervals of CARs from +1 to +10 for the *WT, MCD1-AID, DE, ECO1-AID, eco1*Δ *wpl1*Δ, *DE eco1*Δ, *wpl1*Δ strains listed in A. Numbers in the corners represent the fold-change of the signal enrichment of the center pixel over the indicated corner pixels.

**Fig. S6.**
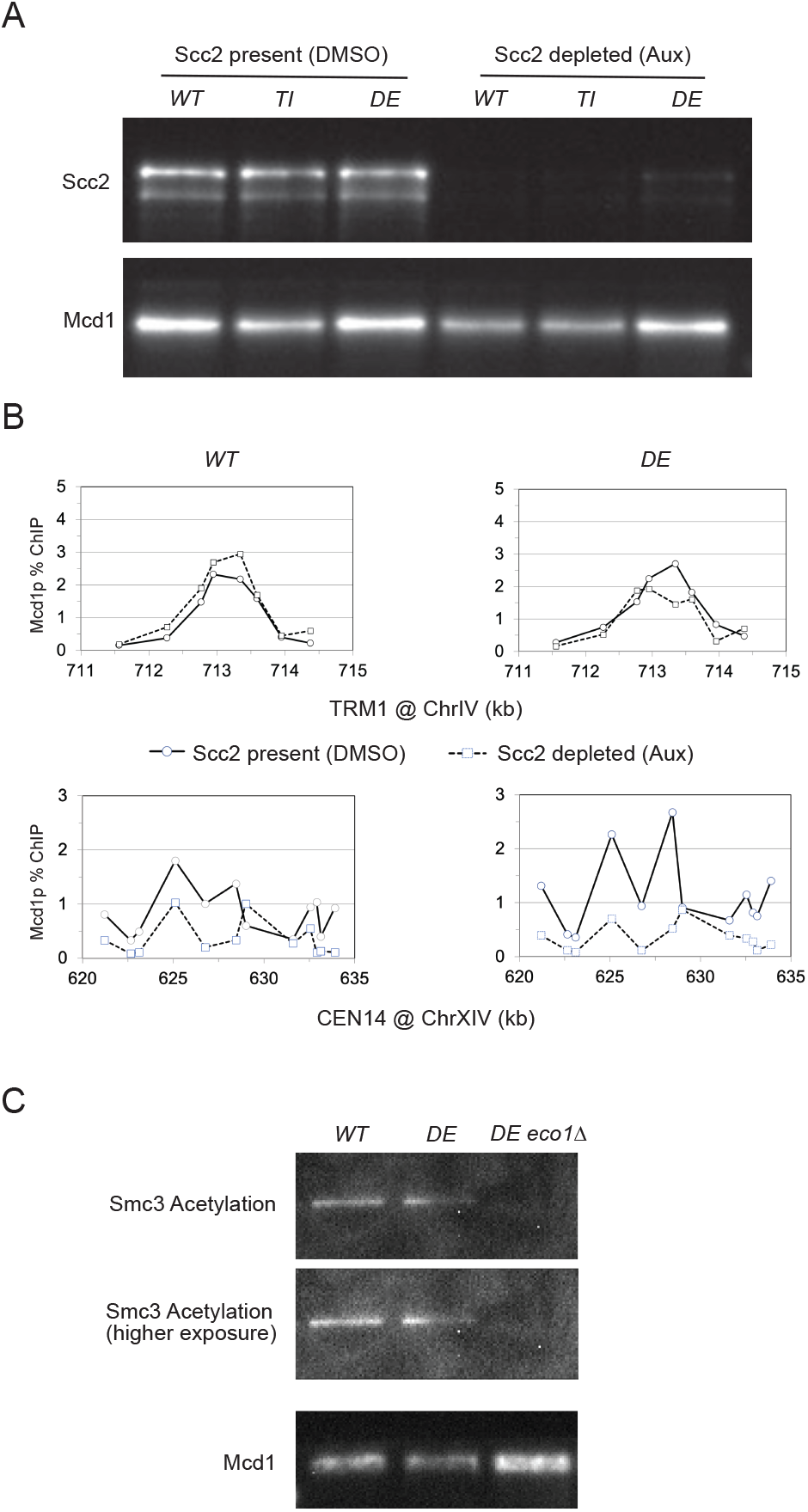
Cohesin smc1-D1164E mutant does not affect cohesin binding stability or acetylation. **(A)** Scc2p is depleted efficiently by auxin. Western blot analysis for Scc2 and Mcd1 using extracts from *WT* cells, *TI* cells, and *DE* cells treated with auxin (Aux) or control (DMSO). **(B)** Cohesin stability on chromatin is not affected by the smc1-D1164E mutation. ChIP qPCR for Mcd1p was performed on wild-type (*WT*) and smc1-D1164E (*DE*) cells, arrested in mitosis and subsequently depleted for Scc2 with auxin (Aux, circles with solid line) or control (DMSO, squares with dotted line). A representative CAR (TRM1 on top) and a centromere (CEN14 on bottom) were probed. **(C)** Smc3 acetylation levels are not affected by smc1-D1164E mutant. Western blot analysis for Smc3 acetylation and Mcd1 using extracts from *WT* cells, *DE* cells, and *DE eco1*Δ cells.

**Supplemental Table 1.**
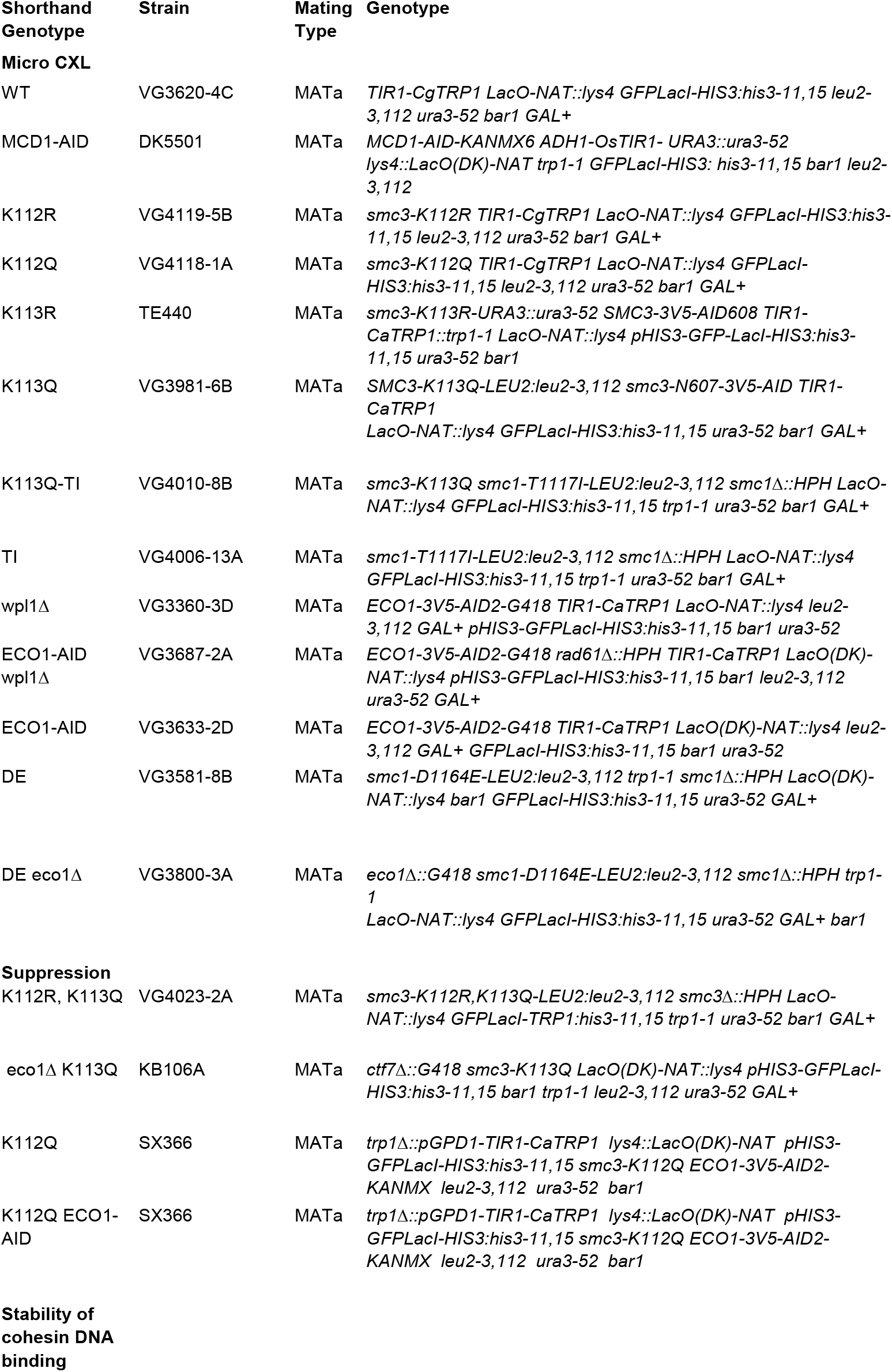
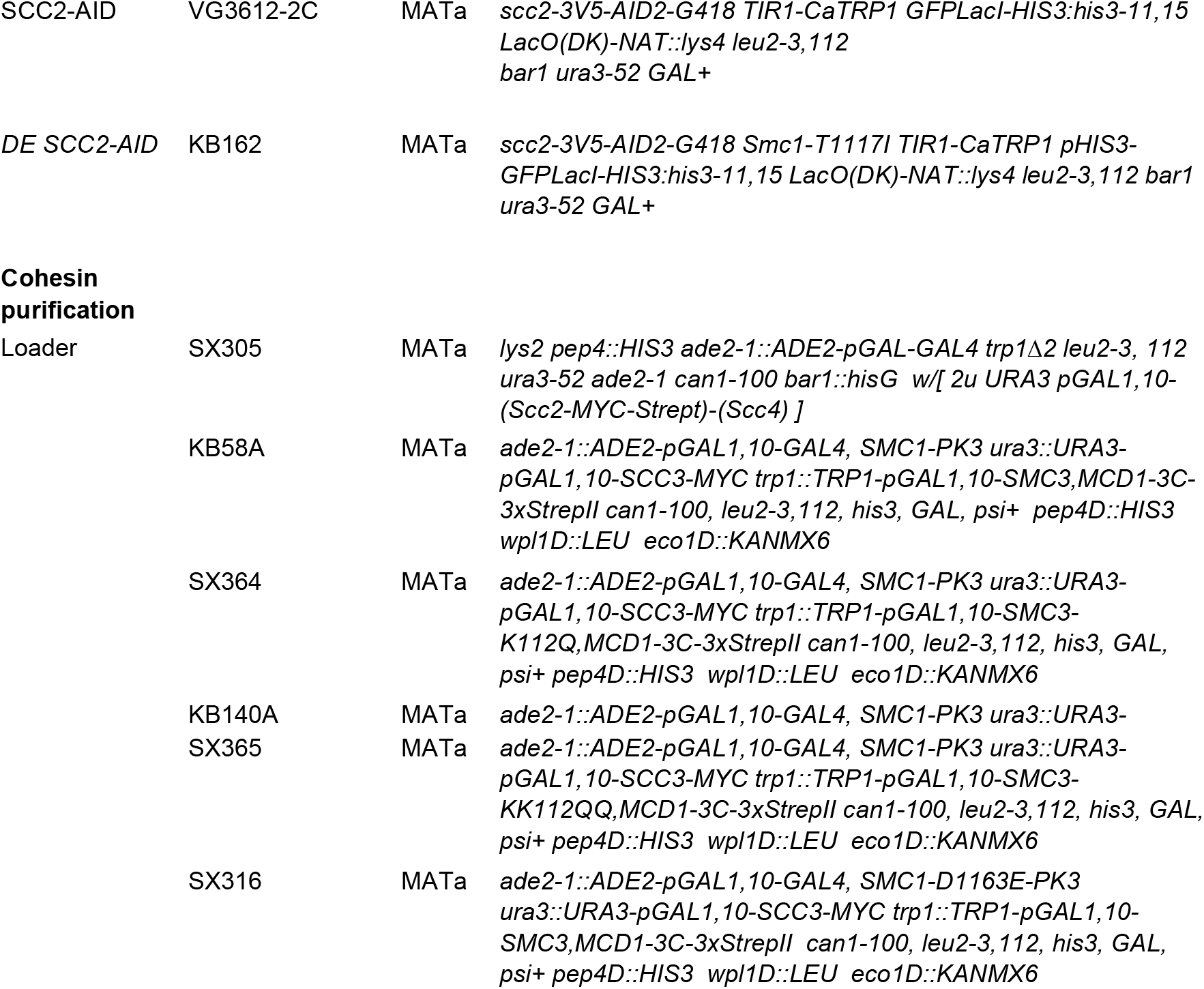

## References

1. Hoencamp, C., and Rowland, B.D. (2023). Genome control by SMC complexes. Nat. Rev. Mol. Cell Biol. 24, 633–650.

2. Carretero, M., Remeseiro, S., and Losada, A. (2010). Cohesin ties up the genome. Curr. Opin. Cell Biol. 22, 781–787.

3. Onn, I., Heidinger-Pauli, J.M., Guacci, V., Unal, E., and Koshland, D.E. (2008). Sister chromatid cohesion: a simple concept with a complex reality. Annu. Rev. Cell Dev. Biol. 24, 105–129.

4. Ganji, M., Shaltiel, I.A., Bisht, S., Kim, E., Kalichava, A., Haering, C.H., and Dekker, C. (2018). Real-time imaging of DNA loop extrusion by condensin. Science 360, 102–105.

5. Kim, Y., Shi, Z., Zhang, H., Finkelstein, I.J., and Yu, H. (2019). Human cohesin compacts DNA by loop extrusion. -PubMed -NCBI. Science 47, eaaz4475–1349.

6. Davidson, I.F., Bauer, B., Goetz, D., Tang, W., Wutz, G., and Peters, J.-M. (2019). DNA loop extrusion by human cohesin. Science 366, 1338–1345.

7. Rao, S.S.P., Huntley, M.H., Durand, N.C., Stamenova, E.K., Bochkov, I.D., Robinson, J.T., Sanborn, A.L., Machol, I., Omer, A.D., Lander, E.S., et al. (2014). A 3D map of the human genome at kilobase resolution reveals principles of chromatin looping. Cell 159, 1665– 1680.

8. Fudenberg, G., Imakaev, M., Lu, C., Goloborodko, A., Abdennur, N., and Mirny, L.A. (2016). Formation of Chromosomal Domains by Loop Extrusion. Cell Rep. 15, 2038–2049.

9. Nora, E.P., Goloborodko, A., Valton, A.-L., Gibcus, J.H., Uebersohn, A., Abdennur, N., Dekker, J., Mirny, L.A., and Bruneau, B.G. (2017). Targeted Degradation of CTCF Decouples Local Insulation of Chromosome Domains from Genomic Compartmentalization. Cell 169, 930–944.e22.

10. Wutz, G., Várnai, C., Nagasaka, K., Cisneros, D.A., Stocsits, R.R., Tang, W., Schoenfelder, S., Jessberger, G., Muhar, M., Hossain, M.J., et al. (2017). Topologically associating domains and chromatin loops depend on cohesin and are regulated by CTCF, WAPL, and PDS5 proteins. EMBO J. 36, 3573–3599.

11. Costantino, L., Hsieh, T.-H.S., Lamothe, R., Darzacq, X., and Koshland, D. (2020). Cohesin residency determines chromatin loop patterns. Elife 9. 10.7554/eLife.59889.

12. Weitzer, S., Lehane, C., and Uhlmann, F. (2003). A model for ATP hydrolysis-dependent binding of cohesin to DNA. Curr. Biol. 13, 1930–1940.

13. Arumugam, P., Gruber, S., Tanaka, K., Haering, C.H., Mechtler, K., and Nasmyth, K. (2003). ATP hydrolysis is required for cohesin’s association with chromosomes. Curr. Biol. 13, 1941–1953.

14. Unal, E., Heidinger-Pauli, J.M., Kim, W., Guacci, V., Onn, I., Gygi, S.P., and Koshland, D.E. (2008). A molecular determinant for the establishment of sister chromatid cohesion. Science 321, 566–569.

15. Rolef Ben-Shahar, T., Heeger, S., Lehane, C., East, P., Flynn, H., Skehel, M., and Uhlmann, F. (2008). Eco1-dependent cohesin acetylation during establishment of sister chromatid cohesion. Science 321, 563–566.

16. Haering, C.H., Löwe, J., Hochwagen, A., and Nasmyth, K. (2002). Molecular architecture of SMC proteins and the yeast cohesin complex. Mol. Cell 9, 773– 788.

17. Skibbens, R.V., Corson, L.B., Koshland, D., and Hieter, P. (1999). Ctf7p is essential for sister chromatid cohesion and links mitotic chromosome structure to the DNA replication machinery. Genes Dev. 13, 307–319.

18. Tóth, A., Ciosk, R., Uhlmann, F., Galova, M., Schleiffer, A., and Nasmyth, K. (1999). Yeast cohesin complex requires a conserved protein, Eco1p(Ctf7), to establish cohesion between sister chromatids during DNA replication. Genes Dev. 13, 320–333.

19. Zhang, J., Shi, X., Li, Y., Kim, B.-J., Jia, J., Huang, Z., Yang, T., Fu, X., Jung, S.Y., Wang, Y., et al. (2008). Acetylation of Smc3 by Eco1 is required for S phase sister chromatid cohesion in both human and yeast. Mol. Cell 31, 143–151.

20. Chan, K.-L., Roig, M.B., Hu, B., Beckouët, F., Metson, J., and Nasmyth, K. (2012). Cohesin’s DNA exit gate is distinct from its entrance gate and is regulated by acetylation. Cell 150, 961–974.

21. Çamdere, G., Guacci, V., Stricklin, J., and Koshland, D. (2015). The ATPases of cohesin interface with regulators to modulate cohesin-mediated DNA tethering. Elife 4, e11315.

22. Bastié, N., Chapard, C., Dauban, L., Gadal, O., Beckouët, F., and Koszul, R. (2022). Smc3 acetylation, Pds5 and Scc2 control the translocase activity that establishes cohesin-dependent chromatin loops. Nat. Struct. Mol. Biol. 29, 575–585.

23. van Ruiten, M.S., van Gent, D., Sedeño Cacciatore, Á., Fauster, A., Willems, L., Hekkelman, M.L., Hoekman, L., Altelaar, M., Haarhuis, J.H.I., Brummelkamp, T.R., et al. (2022). The cohesin acetylation cycle controls chromatin loop length through a PDS5A brake mechanism. Nat. Struct. Mol. Biol. 29, 586–591.

24. Murayama, Y., and Uhlmann, F. (2015). DNA Entry into and Exit out of the Cohesin Ring by an Interlocking Gate Mechanism. Cell 163, 1628–1640.

25. Boardman, K., Xiang, S., Chatterjee, F., Mbonu, U., Guacci, V., and Koshland, D. (2023). A model for Scc2p stimulation of cohesin’s ATPase and its inhibition by acetylation of Smc3p. Genes Dev. 37, 277–290.

26. Ciosk, R., Shirayama, M., Shevchenko, A., Tanaka, T., Toth, A., Shevchenko, A., and Nasmyth, K. (2000). Cohesin’s binding to chromosomes depends on a separate complex consisting of Scc2 and Scc4 proteins. Mol. Cell 5, 243–254.

27. Hsieh, T.-H.S., Weiner, A., Lajoie, B., Dekker, J., Friedman, N., and Rando, O.J. (2015). Mapping Nucleosome Resolution Chromosome Folding in Yeast by Micro-C. Cell 162, 108–119.

28. van Ruiten, M.S., and Rowland, B.D. (2018). SMC Complexes: Universal DNA Looping Machines with Distinct Regulators. Trends Genet. 34, 477–487.

29. Mitter, M., Gasser, C., Takacs, Z., Langer, C.C.H., Tang, W., Jessberger, G., Beales, C.T., Neuner, E., Ameres, S.L., Peters, J.-M., et al. (2020). Sister-chromatid-sensitive Hi-C reveals the conformation of replicated human chromosomes. bioRxiv, 2020.03.10.978148. 10.1101/2020.03.10.978148.

30. Oomen, M.E., Hedger, A.K., Watts, J.K., and Dekker, J. (2020). Detecting chromatin interactions between and along sister chromatids with SisterC. Nat. Methods 17, 1002–1009.

31. Stead, K., Aguilar, C., Hartman, T., Drexel, M., Meluh, P., and Guacci, V. (2003). Pds5p regulates the maintenance of sister chromatid cohesion and is sumoylated to promote the dissolution of cohesion. J. Cell Biol. 163, 729–741.

32. Murayama, Y., and Uhlmann, F. (2014). Biochemical reconstitution of topological DNA binding by the cohesin ring. Nature 505, 367–371.

33. Guacci, V., and Koshland, D. (2012). Cohesin-independent segregation of sister chromatids in budding yeast. Mol. Biol. Cell 23, 729–739.

34. Eng, T., Guacci, V., and Koshland, D. (2014). ROCC, a conserved region in cohesin’s Mcd1 subunit, is essential for the proper regulation of the maintenance of cohesion and establishment of condensation. Mol. Biol. Cell 25, 2351–2364.

35. Birot, A., Eguienta, K., Vazquez, S., Claverol, S., Bonneu, M., Ekwall, K., Javerzat, J.-P., and Vaur, S. (2017). A second Wpl1 anti-cohesion pathway requires dephosphorylation of fission yeast kleisin Rad21 by PP4. - PubMed - NCBI. EMBO J., e201696050.

36. Nagasaka, K., Davidson, I.F., Stocsits, R.R., Tang, W., Wutz, G., Batty, P., Panarotto, M., Litos, G., Schleiffer, A., Gerlich, D.W., et al. (2023). Cohesin mediates DNA loop extrusion and sister chromatid cohesion by distinct mechanisms. Mol. Cell 83, 3049– 3063.e6.

37. Rowland, B.D., Roig, M.B., Nishino, T., Kurze, A., Uluocak, P., Mishra, A., Beckouët, F., Underwood, P., Metson, J., Imre, R., et al. (2009). Building sister chromatid cohesion: smc3 acetylation counteracts an antiestablishment activity. Mol. Cell 33, 763–774.

38. Ladurner, R., Kreidl, E., Ivanov, M.P., Ekker, H., Idarraga-Amado, M.H., Busslinger, G.A., Wutz, G., Cisneros, D.A., and Peters, J.-M. (2016). Sororin actively maintains sister chromatid cohesion. EMBO J. 35, 635– 653.

39. Guérin, T.M., Barrington, C., Pobegalov, G., Molodtsov, M.I., and Uhlmann, F. (2024). An extrinsic motor directs chromatin loop formation by cohesin. EMBO J. 10.1038/s44318-024-00202-5.

40. Chan, K.-L., Gligoris, T., Upcher, W., Kato, Y., Shirahige, K., Nasmyth, K., and Beckouët, F. (2013). Pds5 promotes and protects cohesin acetylation. Proc. Natl. Acad. Sci. U. S. A. 110, 13020–13025.

41. Yu, D., Chen, G., Wang, Y., Wang, Y., Lin, R., Liu, N., Zhu, P., Liu, H., Hu, T., Feng, R., et al. (2022). Regulation of cohesin-mediated chromosome folding by PDS5 in mammals. EMBO Rep. 23, e54853.

## Methods Reference

1. Servant, N., Varoquaux, N., Lajoie, B.R., Viara, E., Chen, C.J., Vert, J.P., Heard, E., Dekker, J. and Barillot, E., 2015. HiC-Pro: an optimized and flexible pipeline for Hi-C data processing. Genome biology, 16(1), p.259.

2. Langmead, B. and Salzberg, S.L., 2012. Fast gapped-read alignment with Bowtie 2. Nature methods, 9(4), pp.357–359.

3. Abdennur, N. and Mirny, L.A., 2020. Cooler: scalable storage for Hi-C data and other genomically labeled arrays. Bioinformatics, 36(1), pp.311–316.

4. Durand, N.C., Shamim, M.S., Machol, I., Rao, S.S., Huntley, M.H., Lander, E.S. and Aiden, E.L., 2016. Juicer provides a one-click system for analyzing loop-resolution Hi-C experiments. Cell systems, 3(1), pp.95–98.

5. Imakaev, M., Fudenberg, G., McCord, R.P., Naumova, N., Goloborodko, A., Lajoie, B.R., Dekker, J. and Mirny, L.A., 2012. Iterative correction of Hi-C data reveals hallmarks of chromosome organization. Nature methods, 9(10), pp.999–1003.

6. Knight, P.A. and Ruiz, D., 2013. A fast algorithm for matrix balancing. IMA Journal of Numerical Analysis, 33(3), pp.1029–1047.

7. Costantino, L., Hsieh, T.H.S., Lamothe, R., Darzacq, X. and Koshland, D., 2020. Cohesin residency determines chromatin loop patterns. Elife, 9, p.e59889.

8. Bastié, N., Chapard, C., Dauban, L., Gadal, O., Beckouet, F. and Koszul, R., 2022. Smc3 acetylation, Pds5 and Scc2 control the translocase activity that establishes cohesin-dependent chromatin loops. Nature structural & molecular biology, 29(6), pp.575–585.

9. Hunter, J.D., 2007. Matplotlib: A 2D graphics environment. Computing in science & engineering, 9(03), pp.90–95.

10. Open2C, Abdennur, N., Abraham, S., Fudenberg, G., Flyamer, I.M., Galitsyna, A.A., Goloborodko, A., Imakaev, M., Oksuz, B.A., Venev, S.V. and Xiao, Y., 2024. Cooltools: enabling high-resolution Hi-C analysis in Python. PLOS Computational Biology, 20(5), p.e1012067

11. Hsieh, T.H.S., Weiner, A., Lajoie, B., Dekker, J., Friedman, N. and Rando, O.J., 2015. Mapping nucleosome resolution chromosome folding in yeast by micro-C. Cell, 162(1), pp.108–119.

12. Rao, S.S., Huntley, M.H., Durand, N.C., Stamenova, E.K., Bochkov, I.D., Robinson, J.T., Sanborn, A.L., Machol, I., Omer, A.D., Lander, E.S. and Aiden, E.L., 2014. A 3D map of the human genome at kilobase resolution reveals principles of chromatin looping. Cell, 159(7), pp.1665–1680.

13. Flyamer, I.M., Illingworth, R.S. and Bickmore, W.A., 2020. Coolpup. py: versatile pile-up analysis of Hi-C data. Bioinformatics, 36(10), pp.2980–2985.

14. Zhang, Y., Liu, T., Meyer, C.A., Eeckhoute, J., Johnson, D.S., Bernstein, B.E., Nusbaum, C., Myers, R.M., Brown, M., Li, W. and Liu, X.S., 2008. Model-based analysis of ChIP-Seq (MACS). Genome biology, 9(9), p.R137.

15. Li, Q., Brown, J.B., Huang, H. and Bickel, P.J., 2011. Measuring reproducibility of high-throughput experiments.

